# Learning Edge Rewiring in EMT From Single Cell Data

**DOI:** 10.1101/155028

**Authors:** Smita Krishnaswamy, Nevena Zivanovic, Roshan Sharma, Dana Pe’er, Bernd Bodenmiller

## Abstract

Cellular regulatory networks are not static, but continuously reconfigure in response to stimuli via alterations in gene expression and protein confirmations. However, typical computational approaches treat them as static interaction networks derived from a single experimental time point. Here, we provide a method for learning the dynamic modulation, or *rewiring* of pairwise relationships (edges) from a static single-cell data. We use the epithelial-to-mesenchymal transition (EMT) in murine breast cancer cells as a model system, and measure mass cytometry data three days after induction of the transition by TGFβ. We take advantage of transitional rate variability between cells in the data by deriving a pseudo-time EMT trajectory. Then we propose methods for visualizing and quantifying time-varying edge behavior over the trajectory and use these methods: TIDES (Trajectory Imputed DREMI scores), and measure of edge dynamism (3DDREMI) to predict and validate the effect of drug perturbations on EMT.

## Introduction

Different cell types exhibit distinct responses to environmental cues, resulting in changes in cellular state. Responses to environmental cues play a key role in development, cellular differentiation and fate. For instance, growth factors like TGFβ [1] guide the patterning of tissues during embryogenesis [2]. In hematopoiesis growth factors such as GM-CSF can drive the development of neutrophils, monocytes and macrophages [3].

During wound healing, TGFβ can induce trans-differentiation from fibroblasts to myofibroblasts. During cell state transitions, configurations of signaling and transcriptional networks rewire, often causing cells to interpret external signals differently and to enable new cellular functions. Our goal is to develop computational methods that infer rewiring of cellular signaling and regulatory networks during state transitions. Further, our methods should facilitate the identification of critical events that are necessary for the transition.

We use the EMT as a model system because it is a substantial change in cell state, that can be generated via a controllable means in model systems. This state transition is important in processes such as embryogenesis, wound healing and some aspects of cancer [4-6]. EMT can be initiated by an external TGFβ signal, resulting in signaling and transcriptional network activation, followed by behavioral and morphological changes. TGF-β-induced EMT is thought to involve among others the SMAD, MAPK and AKT pathways, which activate multiple transcription factors such as Snail, Slug, Twist and Zeb and in turn their targets [7, 8]. The induction of these factors causes cells to lose epithelial characteristics such as the cell-to-cell adhesion and polarity, and acquire mesenchymal characteristics such as migratory competence and spindle-like shape. We study EMT using a mouse breast-cancer cell line [9, 10], and use mass-cytometry to follow the transition of cell state starting with the induction by TGFβ.

Signaling networks can be learned from high dimensional single cell data. For example, Sachs *et.al.* [11] used Bayesian networks to qualitatively learn signaling influences using flow cytometry measurements. Krishnaswamy *et.al.* [12] inferred a quantitative model of signaling interactions from mass cytometry data using a mutual information-based metric known as DREMI. However, most single cell datasets include only a single time point and hence most network-learning methods infer static versions of a time-varying phenomenon (a single network model). By contrast, we recognize that signaling networks are dynamic and reconfigurable entities and that a dynamic view is essential for describing state transition processes like EMT. To support this view, we propose a method that learns dynamic changes in signaling as a continuous process *from* static snapshots of data. This enables us to see how the signaling network is changing over a transition like EMT.

In EMT we observe considerable cell-to-cell variability in the rate at which cells transition. To glean dynamics from a single time point, we order cells onto a one-dimensional trajectory that approximates the EMT progression [13, 14]. We developed new statistical methods, extending on our previous work [12] to model how signaling networks change continuously through EMT, treating relationships between molecules as a time-varying signaling edge. We validate our rewiring assessment using acute inhibitions that supports our inferred changes in signaling relationships. Our continuous view of rewiring also gives us insight into critical edges that may be essential for driving the transition. We find that inhibiting molecules that participate in such critical edges modulate the transition.

## Results

### Measuring signaling during TGFβ-induced EMT

To study the signaling network and phenotypic changes during EMT (Figure 1A), we used Py2T murine breast cancer cells following chronic exposure to TGFβ [15] (Figure 1B). Cells were sampled daily in biological triplicate over a four-day TGFβ time course. We used mass cytometry [16] to assay transcription factors and phosphorylation site abundance regulating signaling protein activity in single cells. A total of 32 markers were simultaneously measured, including three surface markers and 29 intracellular markers. The markers were chosen to assess epithelial (high expression of E-cadherin and CD24) and mesenchymal (high expression of Vimentin and CD44) states, signaling activity of the SMAD, AKT, MAPK, WNT and NFκB pathways, EMT transcription factors, cell cycle, and apoptosis (Supplementary Table 1).

**Figure 1.**
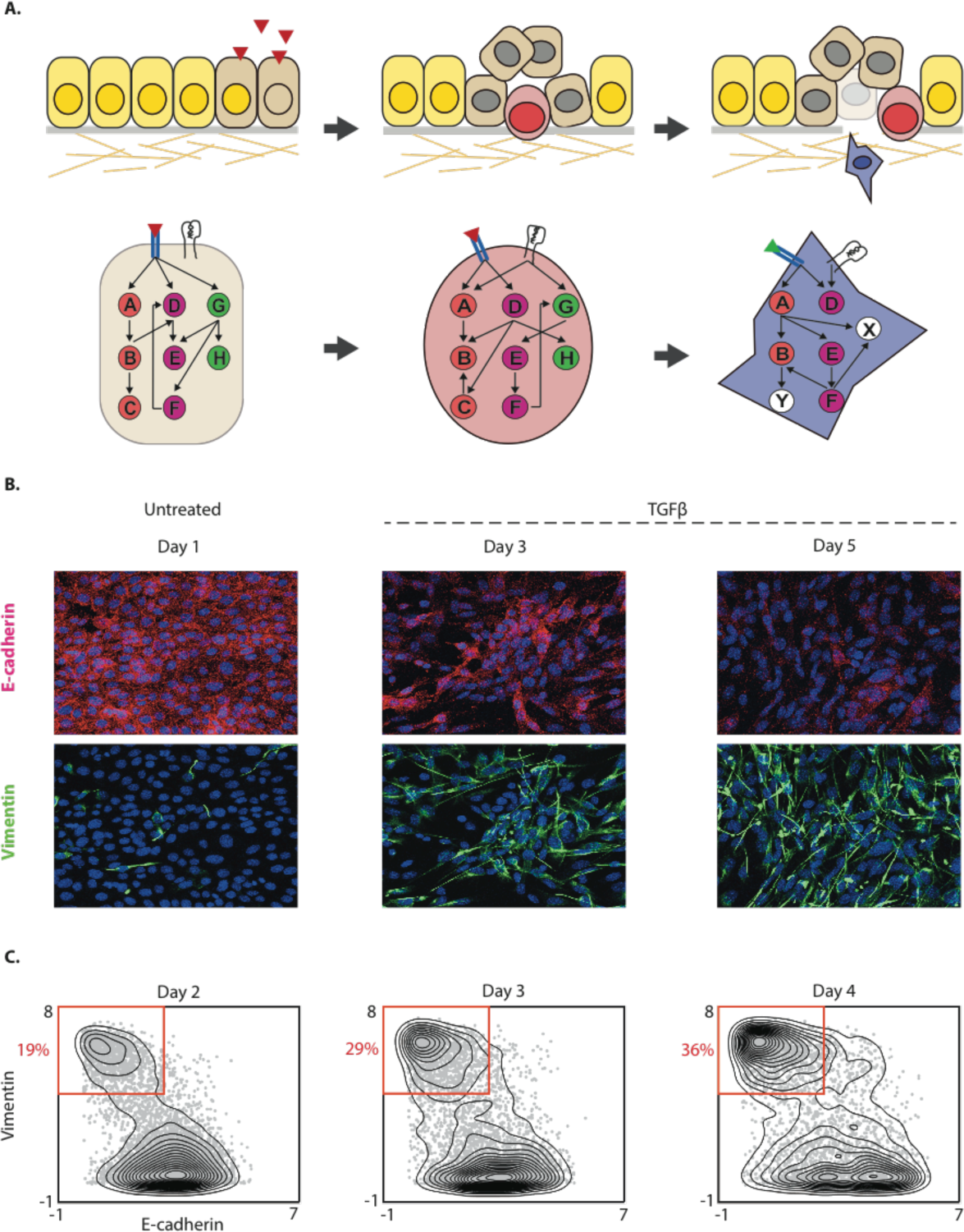
Rewiring during phenotypic change in EMT: (A) Conceptual diagram of rewiring as cells undergo EMT. (B) Immunofluorescence images of Py2T cells stained for canonical markers E-cadherin (in red) and Vimentin (in green) are shown after 1, 3 and 5 days of 4ng/ml TGFβ stimulation. Three days after TGFβ treatment we find both cells that express E-cadherin and cells that express Vimentin. (C) Contour plots of Vimentin and E-cadherin following 2-4 days of TGFβ exposure show a shift in density from epithelial to mesenchymal with 19%, 29% and 36% of cells in the mesenchymal phase respectively. The data is arcsinh transformed with a cofactor of 5. The plots show a continuum of intermediate cell states indicating that EMT is a rate-heterogeneous process.

Starting on day two, we observed cells ranging from the epithelial to the mesenchymal state. Days two, three and four had 19%, 29% and 36% of cells in the mesenchymal state (Figure 1C, Supplementary Figures 1A-B). The epithelial cells showed high levels of E-cadherin and CD24. Cells labeled as mesenchymal recapitulated mesenchymal characteristics, including loss of E-cadherin and gain of Vimentin. The transitioning cells exhibited intermediate marker expression that shared both epithelial and mesenchymal characteristics, based on the expression of E-cadherin, CD24, CD44 and Vimentin (Supplementary Figures 2A-E). Taken together, we observed a continuum of cells from the epithelial to the mesenchymal state, underlining the transitional character of the system, that either implies a high-degree of variability in the rate of transition or in commitment of a given cell to the EMT state. Therefore, rather than treating EMT as a two state-system, in all subsequent analyses we treated the heterogeneous population of cells as a continual trajectory and ordered cells along a pseudo-time axis of EMT progression, inferred using the Wanderlust algorithm [13]. We call the Wanderlust pseudo-time ordering “EMT-time” (Figures 2A-B).

**Figure 2.**
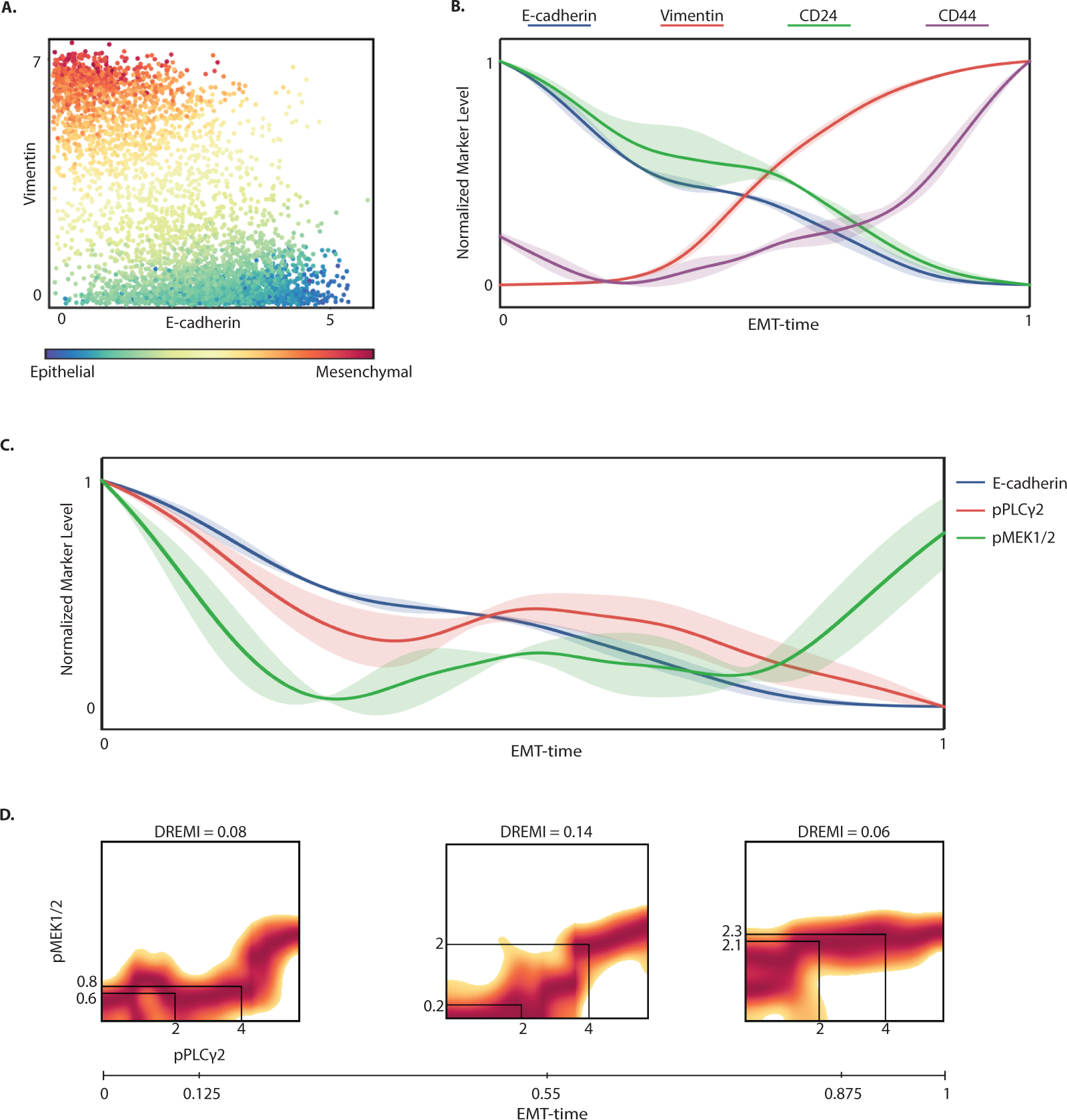
Expression of molecules along EMT-time (wanderlust pseudo-time): (A) Scatterplot where each point represents a Py2T cell collected 3 days after TGFβ stimulation, colored by their Wanderlust-derived pseudo-time label, which we call “EMT-Time” [13]. (B) Smoothed expression levels of E-cadherin, Vimentin, CD24 and CD44 along EMT-time. The EMT-time is normalized to a scale of 0-1, where epithelial cells are near 0 and mesenchymal cells are near 1. Marker levels are also normalized to 0-1 and are smoothed using a sliding-window Gaussian filter. The shaded region around each curve captures 1-standard deviation across replicates, indicating consistency. (C) Smoothed expression levels of signaling markers pPLCγ2 and pΜΕΚ1/2, as well as E-cadherin along EMT-time. (D) DREVI (conditional-Density Rescaled Visualization) [12] plots show the relationship between pPLCγ2 and pMEK1/2 at three different points along EMT-time corresponding to epithelial, transitional and mesenchymal phenotypes. Each DREVI plot illustrates the renormalized conditional density estimate of the abundance of pMEK1/2 given the abundance of pPLCγ2. The red color indicates the conditionally dense regions. The solid black lines indicate that an equal rise in the level of pPLCγ2 results in a higher increase in the abundance of pMEK1/2 during the transitional phase as compared to the epithelial and mesenchymal phase. The strength of the relationship is quantified by DREMI, which computes mutual information on the conditional probability between two molecules. This higher dependency of pMEK1/2 on pPLCγ2 results in a high DREMI score during the transitional phase.

### Extracting an EMT progression from static mass cytometry data via Wanderlust

Given multi-dimensional single cell data, Wanderlust infers a one-dimensional axis of progression and has been shown to accurately recapitulate developmental trajectories [13, 14]. We applied Wanderlust separately to cells from days two, three and four after EMT induction using a subset of the measured markers (Supplementary Table 2). EMT-time recapitulated expected changes: E-cadherin and CD24 showed a monotonic decrease in abundance while Vimentin and CD44 showed a monotonic increase through the transition (Figure 2B and Supplementary Figures 3A-C). The marker expression trend is robust across replicates (mean cross-correlation > 0.87) (Supplementary Figure 3D). Moreover, the inferred trajectories are similar at different days following EMT induction. Supplementary Figure 3E shows that the Wanderlust trajectories are closely correlated between days 2, 3 and 4 (mean cross-correlation > 0.79), suggesting that EMT-time might represent a cell-state that is agnostic to the day of measurement, once the full range of cells from the epithelial to the mesenchymal state are present.

**Figure 3.**
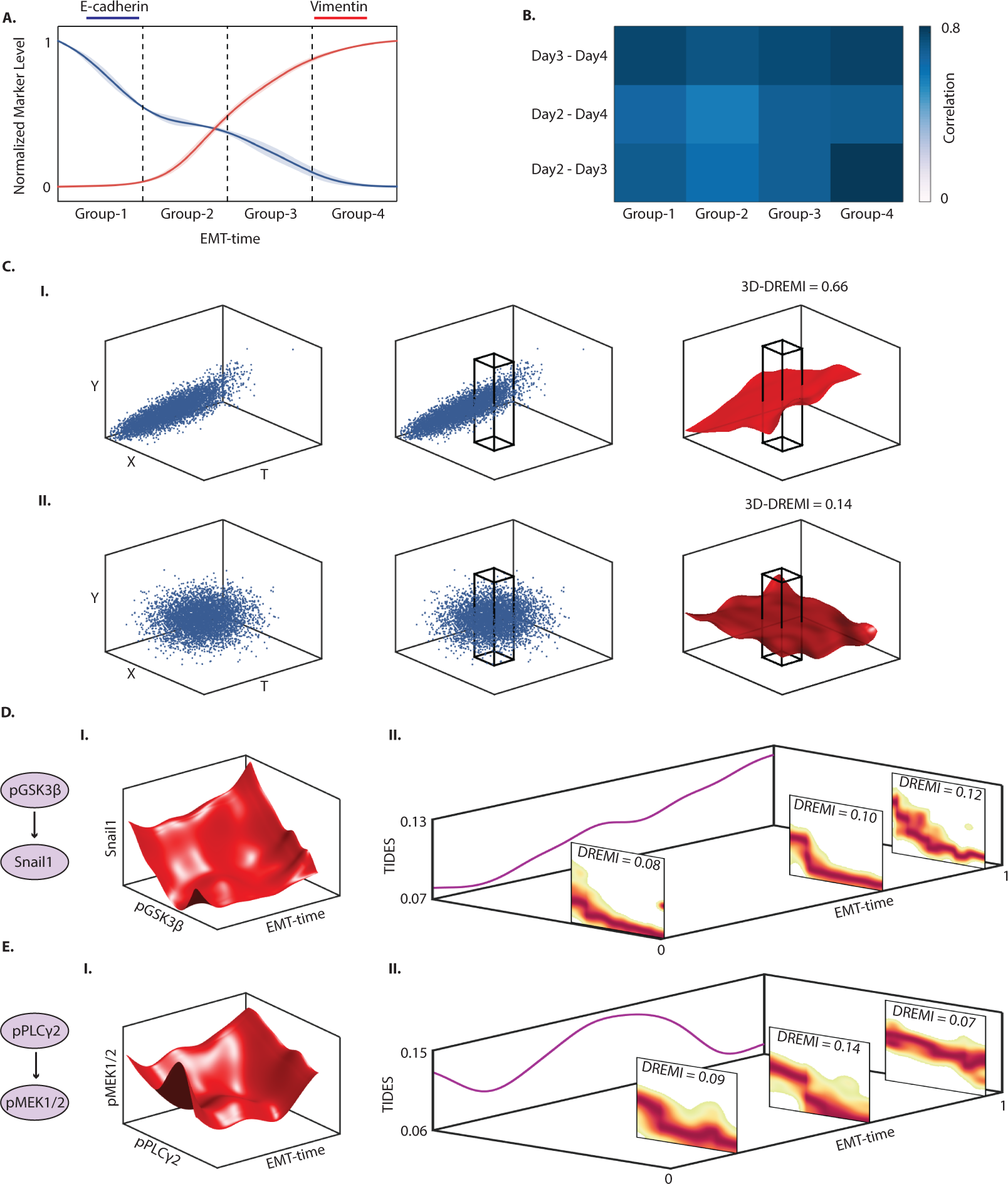
Signaling relationships along EMT-time: (A)-(B) Relationship between signaling molecules is similar across days when controlled for EMT-time. (A) TGFβ-treated cells from Days 2, 3 and 4 are binned into four groups along EMT-time. Expression levels of E-cadherin and Vimentin are shown for reference. (B) Heat map shows the correlation of DREMI scores computed on all pairs of signaling molecules in each group across days. The mean correlation is 0.72. (C) Illustration of computing 3D-DREMI, where each molecule is conditioned by two other molecules. In our work, Y is conditioned on EMT-Time (T) along with the levels of another molecule X. (I) Simulated data depicting strong dependence of Y on both T and X. Across the dynamic range of T and X, Y has a wide range of values. Conditioning Y on both T and X, depicted by the box (middle), substantially decreases the range in Y. The relationship can be seen using a 3D-DREVI plot (right), where the red surface shows the average value of Y conditioned on T and X. Specifying values of T and X provides information on the possible value of Y implying that the relationship has a high 3D-DREMI score. (II) Illustrates a weak relationship with little dependence of Y on X or T. (D) (I) 3D-DREVI between pGSK3β and Snail1 along EMT-time on Day 3 data. (II) The pseudo-dynamics of the relationship between pGSK3β and Snail1 along EMT progression is represented by the TIDES curve (purple curve) which shows the time-varying change in relationship strength (depicted on the Z axis) in the units of DREMI. The 2D-DREVI slices depict the normalized conditional density estimate of the abundance of Snail1 given the abundance of pGSK3β at three specific time-points during EMT. (E) (I) A 3D-DREVI plot of the relationship between pPLCγ2 and pMEK1/2 on Day 3. (II) The TIDES curve and slices of 2D-DREVI along EMT progression show the dynamics of the pPLCγ2->pMEK1/2 relationship, which peaks in strength in an intermediate stage and weakens as the cells complete transition.

### Signaling edges along EMT progression

Using the Wanderlust-derived EMT progression, we studied the *rewiring* or modulations of pairwise relationships i.e., *edges* in the signaling network. Such relationships are statistical dependencies and may or may not reflect direct phosphorylation or other specific biological mechanisms. Studying the dynamics of protein expression and protein phosphorylation levels can tell us which pathways are modulated. Studying an *edge* over time can tell us about how influences between molecules and pathways change. We first compared a canonical edge pPLCγ2-pMEK1/2 between the epithelial, transitional and mesenchymal states using DREMI and DREVI [12] to quantify and visualize edge strength. Figure 2C illustrates how the expression of pMEK1/2 and pPLCγ2 change along EMT-time. Figure 2D shows that the DREVI plot and DREMI scores are substantially different between the states along the progression. The DREMI score between pPLCγ2 and pMEK1/2 increases from 0.08 to 0.14 from the epithelial to transitional state, and subsequently decreases to 0.06 as the cells approach the mesenchymal state (Figure 2D). Thus, the abundance of pPLCγ2 holds more information on the levels of pMEK1/2 in the intermediate state: the same increase in the abundance of pPLCγ2 (from 2 to 4) corresponds to only a small increase in the abundance of pMEK1/2 in epithelial cells (from 0.6 to 0.8) and mesenchymal cells (from 2.1 to 2.3) but a higher increase in transitional cells (from 0.2 to 2).

We sought to confirm whether signaling relationships were more dependent on the actual time point after TGFβ induction of EMT (wall time), or more on a cell’s position in the EMT progression as derived by Wanderlust (EMT-time). The latter is a possibility when different cells progress at individual rates through a fixed EMT program. We binned cells into four stages based on our inferred EMT-time (Figure 3A). For each bin, we computed DREMI scores for all pairs of signaling proteins. We found a high mean correlation of 0.72 between the DREMI scores across days (Figure 3B) when controlled for phase-of-transition (i.e., bins along EMT-time). This result also holds true across various replicates (Supplementary Figure 4A). This result suggests that in our experimental system many signaling relationships are determined by the phase, whereas differences in behavior between time points (wall time) in bulk measurements largely derive from the different proportions of cells in each phase.

Since our data indicates a continuous trajectory with transitional cells between the epithelial and mesenchymal states, we formulated a method to model how relationships between proteins continuously rewire over the course of the EMT progression. We selected Day 3 as a representative sample where cells were relatively uniformly spread throughout the transition. This sets the stage for analysis of protein signaling relationships and their dynamics during the EMT cell-state transition.

### A method to infer rewiring of regulatory edges

To gain a continuous view of edge rewiring (for any particular edge X - Y), we extended DREVI to a 3^rd^ dimension, where the level of the molecule Y is modeled as a function of two parameters: the abundance of the molecule X and EMT-time (T). DREVI is based on the empirical conditional density, estimated directly from the data. As dimensionality increases, data becomes sparser and therefore robust density estimation becomes more challenging. We extended the heat-equation based kernel density estimation [17] used in [12] to higher dimensions (See Methods). We then normalize the density estimate by *two* parent dimensions, rather than one dimension as in [12], to derive the conditional distribution on an X-T plane. We typically visualize a red surface representing the conditionally dense portion of DREVI surface that shows Y’s “typical” behavior for each level of X and point T along EMT-time (Figure 3C(I), right).

Once the ***3D-DREVI*** is computed, we can compute ***3D-DREMI***, measuring the degree of information X *and* T together provide for the value of Y, analogous to 2D-DREMI [12] (see Methods). In Figure 3C(I) the range of Y considerably drops when binned on X and T, thus its 3D-DREMI is high, relative to Figure 3C(II) (low 3D-DREMI) where the range does not change.

While 3D-DREMI provides a general score indicating the degree in which both X and T influence Y, it does not directly address how edge strength changes over time. To derive a quantification of the change in edge strength over the course of the trajectory we introduce a new dynamic measure of dependency that we call *<u>T</u>rajectory <u>I</u>nterpolated <u>D</u>R<u>E</u>MI <u>S</u>cores (TIDES).* A TIDES curve is computed by first computing a 3D conditional density estimate *f*(T, X, Y) where T is EMT-time and X-Y are two molecules whose time varying dependency we intend to assess. Next we compute 2D DREMI in slices along fixed values T (EMT-time) by linearly interpolating the 3D conditional density at small intervals to obtain interpolated 2D DREVI slices and computing the DREMI of these slices (See Figure 3D-E, Methods for details). Thus, the projection of the 3D-conditional density on to a slice allows us to compute the DREMI score between the two markers at any given EMT-time. When taking a causal interpretation of an edge (possibly due to prior knowledge of mechanism), higher DREMI suggests that X exerts a stronger influence on Y. Computing DREMI at each point along EMT-time results in a TIDES curve, which provides a concise, quantitative view describing how pairwise molecular relationships change during the progression.

### A continuous view of rewiring during EMT

*TIDES* allows us to examine how the relationship between two molecules evolves during a state transition. For example, the relationship between signaling molecule GSK3β and the transcription factor Snail1, Figure 3D(I). GSK3β phosphorylates Snail1 at two motifs and is known to inactivate its transcriptional activity and cause protein degradation [7]. However, phosphorylation of GSK3β (pGSK3β) (e.g. through the AKT and PKC pathways [5]) inhibits its activity and therefore pGSK3β is positively correlated with Snail1. Snail1, in turn, modulates genes relevant to EMT and among others activates additional transcription factors [7]. The strength of the relationship between pGSK3β and Snail1 is weak at the beginning of the transition (DREMI = 0.08) and then grows steadily and peaks as the cells are on the verge of completing the transition (DREMI = 0.12), Figure 3D(II). This change is consistent across replicates (Supplementary Figure 4B).

Another example is the edge between phosphorylated PLCγ2 (pPLCγ2) and phosphorylated MEK1/2 (pMEK1/2) shown in Figure 3E. This relationship increases and peaks in strength during the transition (DREMI = 0.14) and wanes again as the transition concludes (DREMI = 0.07). It is known that PLCγ2 enzymes are activated by receptor tyrosine kinases in response to growth factors [18, 19], making PLCγ2 one of the first players that get phosphorylated following growth factor stimulation. pPLCγ2 can then induce the MEK/ERK pathway via PKC [20-22]. In our case, the receptors are the TGFβ receptors (R1 and R2) [23]. We find that this pathway is transmitting the most information during the transitional phase, as indicated by the high DREMI score. This change is consistent across replicates (Supplementary Figure 4C).

In addition to analyzing edges individually, TIDES can also be used to globally understand when there is a high information flow in the entire system. There are points in EMT-time when many signaling molecules pass signal to transcription factors (high DREMI). It can be assumed that such points of high information transfer correspond to critical points, when the system is going through a phase transition. In Figure 4, we combine TIDES scores incoming into the EMT transcription factor Slug, a known core-regulator of EMT [24] which in turn regulates additional EMT transcription factors such as Twist [25]. The average TIDES curve of signaling molecules into Slug shows the scores start rising at around EMT-time 0.25, around when the cell morphology begins to change and hence corresponds to where the transition is beginning. We see a peak towards the end of the transition at around EMT-time 0.8 which might correspond to an additional change in cell state, perhaps when the transition becomes stable (Figure 4). Additionally, we see similar behavior at similar EMT-times for two more EMT transcription factors, Snail1 and Twist (Supplementary Figure 5A-B).

**Figure 4.**
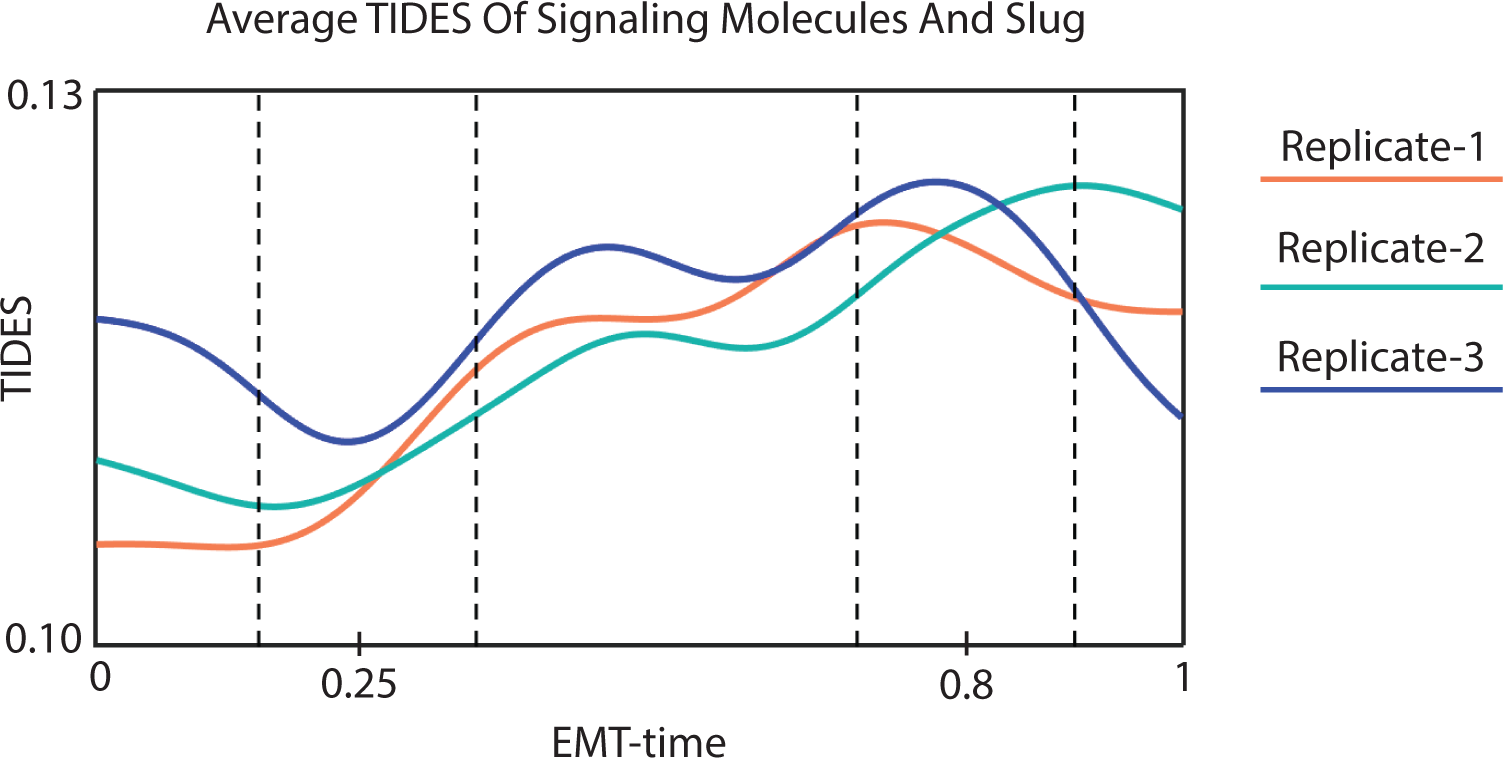
Information transfer during EMT: Average TIDES curve of the relationship between several molecules (pCREB, pSTAT5, pFAK, pMEK1/2, pNFκB, pP38, pAMPK, pAKT, pERK1/2, pGSK3β, pSMAD1/5, pSMAD2/3, β-catenin, CAH IV, pMARCK, pPLCγ2, pS6, pSTAT3) and Slug, across three replicates for Day 3. The curves start rising steadily at near EMT-time = 0.25, and peak near EMT-time = 0.8.

### Validation of TIDES with Acute Inhibitions

Our analysis indicates that the strength of the relationship between signaling molecules changes during the course of EMT. To validate some of the rewiring suggested by TIDES, we performed experiments inhibiting MEK1/2 with the small molecule PD184352. Py2T cells were treated with TGFβ for 3 days, followed by small molecule inhibition of MEK1/2 for 30 minutes prior to cell sampling. Inhibiting the kinase for 30 minutes should accentuate the immediate downstream effects on signaling pathways without substantially altering transcriptional activity, EMT phenotype, or allowing for compensatory effects. Hence, we can directly compare EMT-time of the control and treated condition. We chose to inhibit MEK because it has a potent and specific inhibitor and we were able to measure proximal downstream phosphorylation targets of MEK1/2 (ERK1/2 and P90RSK) by mass cytometry.

For a given edge X-Y, we measured the impact of perturbing X on the abundance of Y. When X-Y represents a causal influence, we expect the impact of this inhibition to correlate with the DREMI at each point in EMT-time, i.e., when DREMI between X and Y is higher, the impact of the inhibition of X on Y is greater and vice versa. We define an *impact curve* as the difference between the abundance of Y along EMT-time under control (no drug-perturbation) and the abundance of Y along EMT-time with drug-perturbation (see Fig. 5A(IV)). We expect regions of high DREMI of X and Y to coincide with the regions of high impact and test this by correlating the TIDES curve against the impact curve, using cross-correlation to match the trajectories (Figure 5A and Methods).

**Figure 5.**
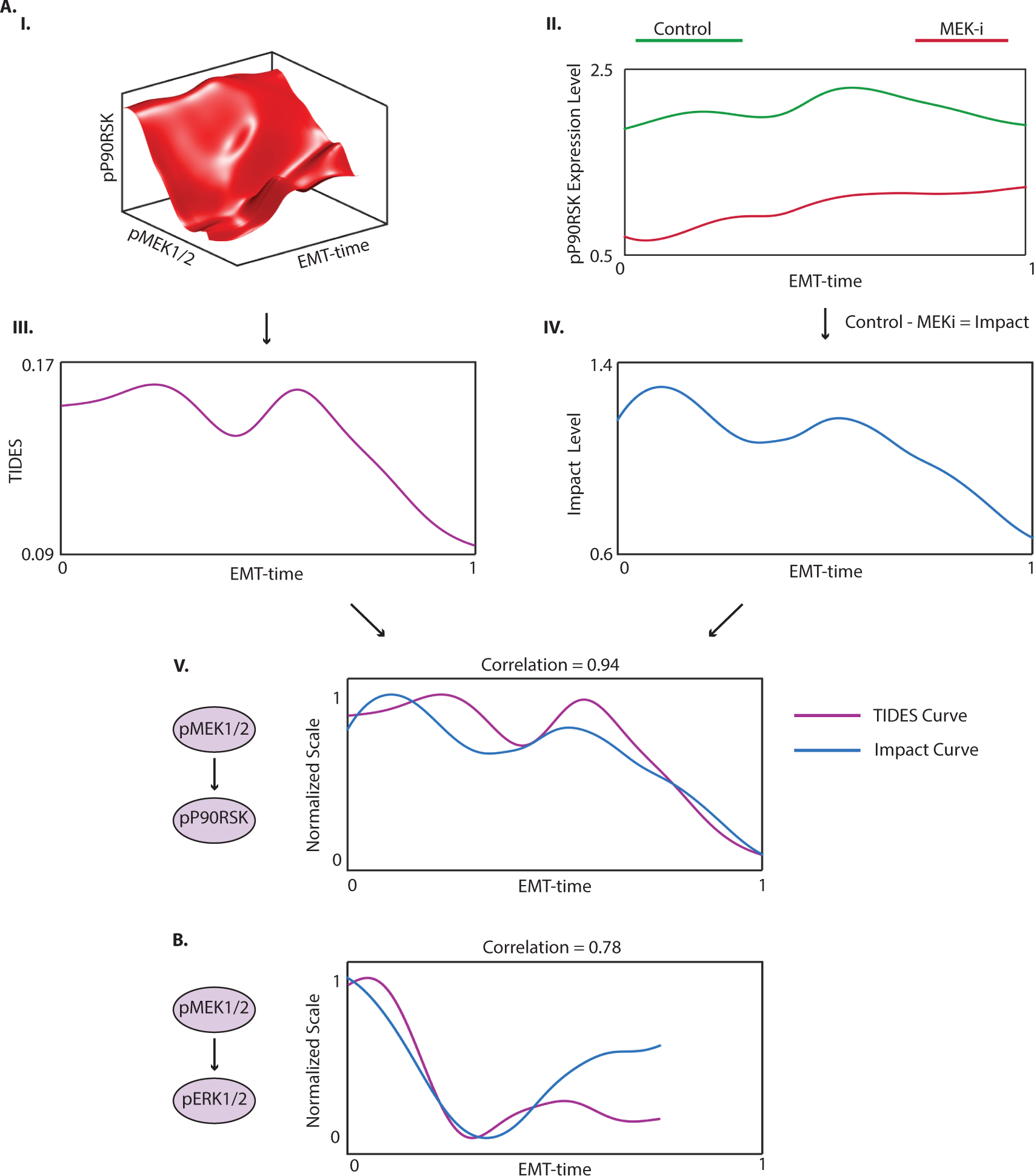
Validation of TIDES via short-term drug inhibition: (A) (I) 3D-DREVI plot shows the typical behavior of pP90RSK given pMEK1/2 and EMT-time. The cells are treated with TGFβ for 3 days. (II) The levels of pP90RSK under control (stimulated with TGFβ) and under MEK-inhibition (TGFβ + MEKi) along EMT-time. As expected, MEK-inhibition substantially reduces the level of pP90RSK as compared to the control. (III) TIDES curve between pMEK1/2 and pP90RSK. (IV) The impact curve, computed as the level of pP90RSK under control minus under MEK-inhibition, shows regions of high effect of MEK-inhibition on pP90RSK along EMT-time. (V) Cross-correlating the curves results in a correlation of 0.94. The depicted curves have been normalized to 0-1 and shifted appropriately based on the lag obtained from cross-correlation (see Methods). (B) Cross-correlating the TIDES curve of pMEK1/2 on pERK1/2 against the impact curve of pERK1/2 under MEK-inhibition also gives a high correlation of 0.78.

We first compared the pP90RSK impact curve along with the pMEK1/2-pP90RSK TIDES curve, Figure 5A. pMEK1/2 is upstream of pP90RSK with pERK1/2 the mediatory kinase that directly phosphorylates p90RSK. We find that the impact curve shows a high cross-correlation of 0.94 with the TIDES curve (Figure 5A(V)), a trend that is repeated across replicates (Supplementary Figure 6A). Note the correlation between the abundance of pP90RSK in control and the TIDES curve is only 0.61 demonstrating that 1) TIDES does not trivially follow levels of the Y-molecule, and 2) that it adds additional predictive value to edge strength (Supplementary Figure 6B). Similarly, the impact curve of pERK1/2 under MEK-inhibition matches the pMEK1/2-pERK1/2 TIDES curve with a cross-correlation of 0.78 (Figure 5B, Supplementary Figure 6C), further validating the approach. The cross-correlation between the pERK1/2-pP90RSK TIDES curve and the impact curve of pP90RSK under MEK-inhibition is also high (0.84 and 0.80 across replicates, Supplementary Figure 6D-E). Thus, we have validated the predictive capability of TIDES in all measured edges downstream of pMEK1/2 in our data.

In the case of MEK-inhibition, the associated relationships were proximal members along a short signaling pathway thus enabling an easy interpretation and thus validation of our approach. Distant relationships can have inputs or convergence from several pathways and therefore the TIDES curve may not accurately match the output. Overall, we find that the TIDES successfully predicts the impact to downstream partners in signaling relationships and can therefore be used to study the time-varying behavior of signaling edges.

### Identification and Validation of Critical Edges in EMT via 3D-DREMI

Next, we wanted to identify edges that are critical drivers for EMT based on rewiring behavior. We hypothesized that driving edges should involve molecules that have a strong dependence on *both* EMT-time, and each other. We wanted to make sure that for a given pair of molecules X-Y and EMT-time (T), X and T *together* provide more information about Y as compared to individually and Y is also highly dependent on both X and T, individually. Therefore, we add together a 3D-DREMI score on (T, X)-Y *and* 2DDREMI on X-Y and T-Y in our panel and sort them by their average score across the three replicates. We find that the top ranking edges are pSMAD2/3 – β-catenin, pAMPK - β-catenin, pERK1/2 - β-catenin, pGSK3β - β-catenin, pGSK3β - pERK1/2 and pMEK1/2 - β-catenin (Supplementary Table 3). This suggests that these molecules, and their corresponding pathways, could be involved in interactions that are strongly regulated during EMT progression. We predict that interrupting these molecules and pathways will have an impact on EMT.

To validate whether our critical edge predictions modulate EMT, we perturb these edges using drug inhibitions and activations. To determine the effect of the modulation on the EMT phenotype, we chronically inhibited/activated the respective molecules and pathways for 5 days while treating the Py2T cells with TGFβ (see Methods). For comparison, cells were only treated with and without TGFβ for the same time. As an additional negative control we used AKT inhibition as an example of a molecule that does not score high in the critical edge list (although typically associated with EMT). We then compare the percentage of cells that transitioned as measured by mass cytometry (Figure 6).

**Figure 6.**
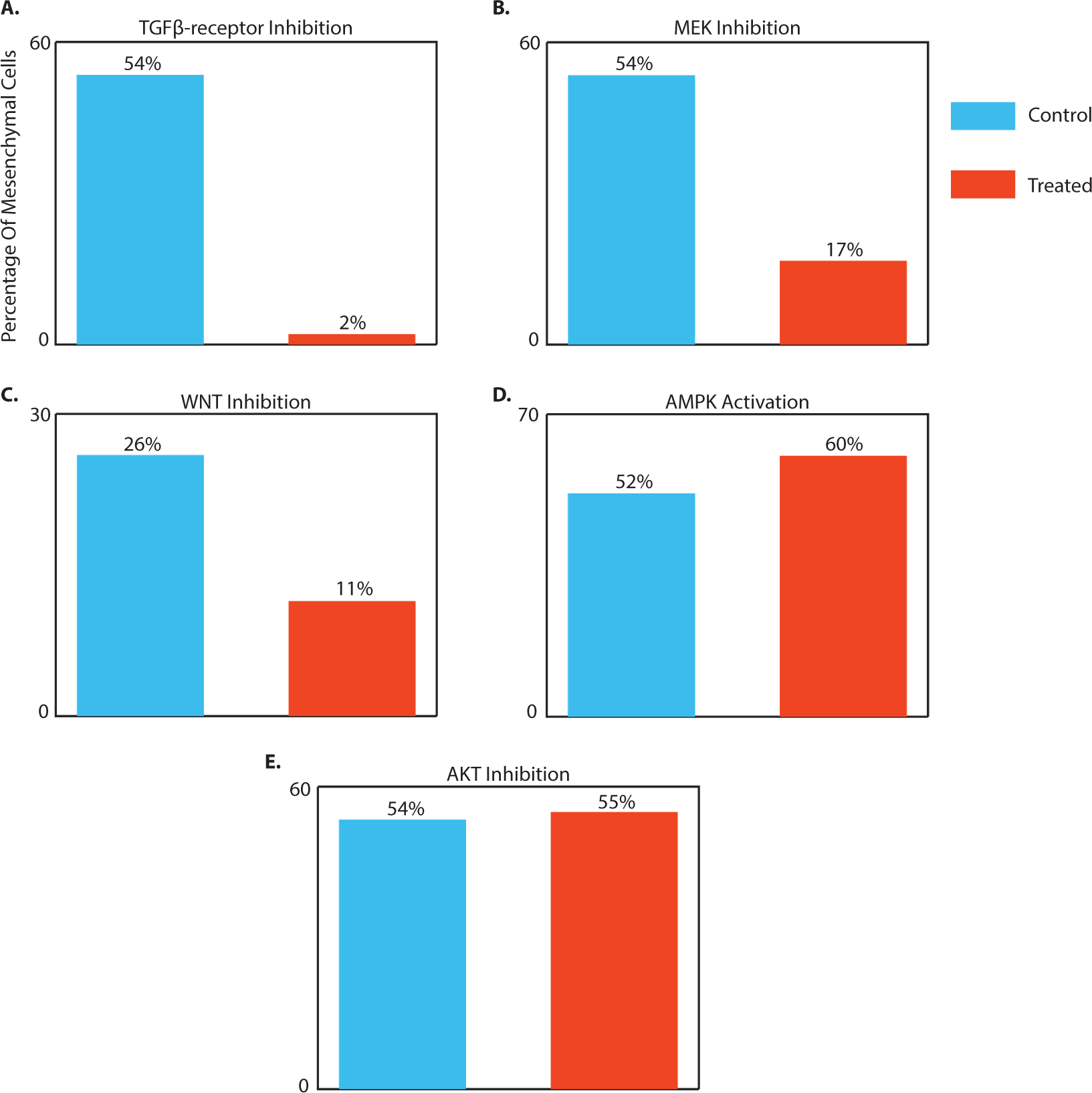
Validation of critical edges for EMT: (A)-(E) Bar plots showing the percentage of mesenchymal cells under control (TGFβ stimulation) and under perturbation of the stated molecule for 5 days. The percentage values measure the impact of the perturbation on EMT. Manual gates were defined to identify mesenchymal cells (see Methods). (A) Inhibition of TGFβ-receptor substantially reduces the fraction of cells completing the EMT transition from 54% under control to 2% following inhibition. (B) MEK inhibition also has a large impact on EMT, under which the fraction of mesenchymal cells drops to 17% from 54% under control. (C) WNT inhibition also causes the fraction of cells completing the transition to drop to 11% from 26% under control. (D) Activating AMPK on the other hand seems to slightly push cells into EMT as the fraction of mesenchymal cells increases from 52% to 60%. (E) AKT inhibition on the other hand has no impact on EMT, fraction of cells completing the transition is 55% compared to 54% under control.

Inhibited molecules/pathways from our predicted critical edges include 1) SMAD2/3, 2) MEK/ERK/MAPK pathway, 3) β-catenin/WNT pathway and 4) AMPK. All results reproduced across biological replicates (Supplementary Figure 7).

1. **SMAD2/3:** Upon TGFβ stimulation, SMAD2/3 is phosphorylated by the TGFβreceptor [26], thus inhibition of the TGFβ-receptor (SB431542) will abrogate SMAD phosphorylation. We find that inhibition of the TGFβ-receptor causes the strongest impact on the progression. The fraction of cells that complete the transition drops to 2% under TGFβ-receptor inhibition as compared to 54% in the control (Figure 6A).
2. **MEK/ERK**: Inhibition of MEK (PD318088) blocks the activity of the MAPK pathway (and therefore also the activity of ERK and P90RSK). Under the MEK-inhibition, the fraction of cells that complete the transition drops to 17% (Figure 6B), less than a 1/3 of the cells that transitioned under control conditions, supporting its role in driving EMT.
3. **β-catenin:** To probe WNT/βcatenin pathway, we use the drug XAV-939 which is known to perturb WNT signaling and cause further β-catenin degradation. Under this inhibition, the fraction of cells that complete the progression drops by 15% (only 11% of the cells transition as compared to 26% in control), see Figure 6C.
4. **AMPK**: For AMPK, we tested an activator rather than an inhibitor, Phenformin. Activation of AMPK slightly increased the percentage of cells that underwent EMT to 60% compared to 52% in control (Figure 6D).

Additionally, we tested the AKT inhibitor (PHT427) as a negative control. AKT has been reported to be an important regulator in EMT for other cell lines such as human squamous carcinoma cells (SCC13 and SCC15) [27] and NMuMMG mammary epithelial cells [28]. Despite the AKT pathway being reported as prominent in the literature [5], we find edges involving AKT to be low in our ranking and indeed we empirically measure that AKT inhibition has little to no impact on EMT (Figure 6E, 55% of cells transition, compared to 54% on control). This result illustrates that EMT is not driven by the same edges across different systems.

In summary, 3D-DREMI successfully predicted the important molecules and pathways involved during EMT, suggesting that the 3D DREMI analysis can be used to generate novel hypotheses on edges that are most relevant for a biological process of interest.

## Discussion

Here, we studied a transition in cell state and how regulatory relations re-wire during this transition. Specifically, we quantified how the strength in relationship between two protein epitopes changed along a developmental trajectory. Importantly, we learned these dynamics from a single time-point of multidimensional single-cell data through a combination of pseudo-time trajectory mapping and dependence tracking along this trajectory. Current high-throughput single-cell techniques offer high-dimensional snapshot measurements of thousands of cells, but without capturing the dynamics. We utilized the fact that cells can exist in various phases of a transitional trajectory due to the variability and asynchrony in transition rates [29]. A major assumption underlying our approach is that while the cells progress through EMT at different rates, they largely do so along a similar path. Therefore, we were able to map the process along a pseudo-time dimension despite the inability to follow a single cell. Once cells were aligned along their position in a pseudo-time trajectory, we tracked how relationships between molecules changed by formulating a dynamic model of edge strength and shape. As such, our approach overcomes the lack of dynamic and continuous single cell data and enables a continuous view on molecular relationships in cell state transition and developmental processes.

All cells in our body contain the same DNA. The difference between cell types lies in the expression and regulation of individual genes and proteins, and the interactions between these proteins. Hence differentiation (or trans-differentiation like in EMT) is essentially a process of gene and protein network rewiring. Thus, treating gene or protein networks as static fundamentally misses key aspects of this rewiring. In our study we analyzed the epithelial-to-mesenchymal transition, which has important roles during development, wound healing, tissue fibrosis and cancer. While this study was limited to this model system, it could be applied to understanding the wiring of the regulatory systems that govern malignant process, opening up exciting possibilities. It is rare that the heterogeneity of response to drug treatment is taken into account, whereas our approach enables the assessment of how a perturbation affects a highly rewired network over the whole continuum of states. We validated our methods by using acute and chronic perturbations in the EMT system. Indeed, times of higher dependence between molecules resulted in a larger impact upon perturbation. Moreover, we confirmed that the perturbation of these highly dynamic molecules, as predicted by our analysis, enabled the identification of nodes that halt EMT. Hence the dynamics can inform us of the key players involved in the developmental (and dynamic) process and aid the selection of drugs that target key factors.

The methods we developed in this paper quantify evolving edge dynamics: specifically we quantified and visualized the time-varying relationship between pairs of molecules and tracked their relationship strength in a continuous manner. Our methods can identify event ordering, peak points of signal transfer, as well as give overall scores for the dynamism of an edge. Further these methods are generic, and can be utilized in any system where single-cell measurements can be used to sample cells that are time-asynchronous with respect to a process and derive the time-dependent dynamics, as well as identification of critically rewired molecules.

Currently single cell technologies are rapidly developing, thus enabling the measurement of ever increasing numbers of molecules of interest. For example, single-cell RNA-sequencing [30, 31] provides a genome wide single cell snap-shot and we could extend our approach to such higher dimensional data types. This could inform us of critical players in transcriptional networks and offer insights into regulatory mechanisms that drive developmental and differentiation processes. However, single-cell RNA sequencing data tends to be sparse, typically capturing only 5-10% of the molecules whereas DREMI and the higher dimensional versions formulated here require sufficient amounts of data to estimate density in the full dynamic range of molecules. However, our approach could be modified to operate under sparse data conditions.

In this study we have formulated a general framework for studying dynamic interactions with static-snapshot data and present the first continuous analysis of the signaling networks controlling the epithelial-mesenchymal transition. We believe that our methods will lay a useful conceptual and quantitative foundation for analyzing relationship dynamics for any type of single cell analysis data.

## METHODS

### Py2T cell culture and stimulation

Py2T cells were obtained from the laboratory of Gerhard Cristofori, University of Basel, Switzerland [1]. Cells were tested for mycoplasma contamination upon arrival and regularly during culturing and before being used for experiments. Cells were cultured at 37 °C in DMEM (D5671, Sigma Aldrich), supplemented with 10% FBS, 2 mM Lglutamine, 100 U/ml penicillin, and 100 μg/ml streptomycin, at 5% CO_2._ For cell passaging, cells were incubated with TrypLE™ Select 10X (Life Technologies) in PBS in a 1:5 ratio (v/v) for 10 minutes at 37°C. For each experiment, cells were seeded at the density of 0.3 million cells per plate (100 mm diameter) and allowed to recover for 36 hours. After reaching 60% confluence, cells were either mock treated or treated with 4ng/ml TGFβ (Human recombinant TGFβ1, Cell Signaling Technologies) for 2, 3 and 4 days. Cell growth media and 4ng/ml TGFβ treatment was renewed every day.

#### Cell harvesting

For cell harvest, cells were washed two times with PBS and incubated with TrypLE™ Select 10X (Life Technologies) in PBS at a 1:5 ratio (v/v) for 10 minutes at 37°C. Following cell detachment, cells were cross-linked by addition of formaldehyde at a final concentration of 1.6% for 10 minutes at room temperature. Cross-linked cells were then centrifuged at 600 × *g* for 5 minutes at 4°C. After aspirating the supernatant, the cell pellet was re-suspended in -20°C methanol to a suspension of 1 million cells/ml and transferred to -80°C for long-term storage.

### Metal-labeled antibodies

Antibodies were obtained in carrier/protein free buffer and labeled with isotopically pure metals (Trace Sciences) using MaxPAR antibody conjugation kit (Fluidigm), according to the manufacturer’s standard protocol. After determining the percent yield by measurement of absorbance at 280 nm, the metal-labeled antibodies were diluted in Candor PBS Antibody Stabilization solution (Candor Bioscience GmbH) for long-term storage at 4°C. Antibodies used in this study are listed in Supplementary Table 1.

### Mass-tag cellular barcoding and antibody staining

Cell samples in methanol were washed three times with Cell Staining Media (CSM, PBS with 0.5% BSA, 0.02% NaN_3_) and once with PBS at 4°C. The cells were then resuspended at 1 million cells/ml in PBS containing barcoding reagents (^102^ Pd,^104^ Pd, ^105^Pd, ^106^Pd,^108^Pd, ^110^Pd,^113^In and^115^In,) each at a final concentration of 100 nM. Cells and barcoding reagent were incubated for 30 minutes at room temperature. Barcoded cells were then washed three times with CSM, pooled and stained with the metal-conjugated antibody mix (Supplementary Table 1) at room temperature for 1 hour. Unbound antibodies were removed by washing cells three times with CSM and once with PBS. For cellular DNA staining, an iridium-containing intercalator (Fluidigm) was diluted to 250 nM in PBS containing 1.6% PFA and added to the cells at 4°C for overnight incubation. Before measurement, the intercalator solution was removed and cells were washed with CSM, PBS, and ddH_2_O. After the last washing step, cells were re-suspended in MilliQ H_2_O to 1 million cells/ml and filtered through a 40-μm strainer.

### Mass cytometry analysis

EQ^™^ Four Element Calibration Beads (Fluidigm) were added to the cell suspension in a 1:10 ratio (v/v). Samples were analyzed on a CyTOF1 (DVS Sciences). The manufacturer’s standard operation procedures were used for acquisition at a cell rate of ~300 cells per second as described in [2]. After the acquisition, all FCS files from the same barcoded sample were concatenated using the Cytobank concatenation tool (http://www.support.cytobank.org/hc/en-us/articles/206336147-FCS-file-concatenationtool). Data were then normalized [3], and bead events were removed. Cell doublet removal and de-barcoding of cells into their corresponding wells was done using a doublet-free filtering scheme and single-cell deconvolution algorithm [4]. Subsequently, data was processed using Cytobank (http://www.cytobank.org/). Additional gating on the DNA channels (^191^Ir and^193^Ir) was used to remove remaining doublets, debris and contaminating particulate.

### Immunofluorescence microscopy analysis

Cells were seeded on 12 mm glass coverslips in 24-well plates. After reaching 60% confluence, cells were treated with TGFβ for 3 and 5 days. The cell growth media containing 4ng/ml TGFβ was replenished once per day. All sample preparation steps were performed at room temperature. Cell samples were cross-linked with 4% paraformaldehyde in PBS for 20 min and permeabilized using 0.1% Triton X-100 in PBS for 3 min. After a blocking step with 0.5% BSA in PBS for 20 min, cell samples were incubated with the primary antibodies (E-Cadherin, Alexa Fluor^®^ 647, 36/E-Cadherin, BD Biosciences; and Vimentin (D21H3) XP^®^ Cell Signaling Technologies) for 1.5 hours, and subsequently incubated for 1 hour with the appropriate fluorophore conjugated secondary antibodies (Alexa Fluor-488). Fluorophore-labeled antibodies were diluted in buffer containing 0.5% BSA in PBS. Nuclei were stained with Hoechst 33258 stain (Sigma Aldrich) diluted in PBS for 3 min. Coverslips were mounted in ProLong^®^ Gold Antifade Mountant (Thermo Fisher Scientific) on microscope slides and imaged with a confocal microscope CLSM SP8 upright Leica. Images were acquired and analyzed using Imaris Software (Bitplane, Switzerland) and the acquisition was performed on the same day to prevent differences due to emission changes of the light sources. In addition, exposure times for a given marker were kept constant for the comparative analysis of each antibody.

### Time course experiment

Mock-treated and TGFβ-treated cells were sampled for measurement after 2, 3 and 4 days. For each condition, three biological replicates were cultured, harvested and analyzed.

### Acute kinase inhibition

After chronic TGFβ stimulation for 3 days, cells were treated with MEK (PD184352) small molecule inhibitors for 30 minutes at a concentration of 10μM and collected in two replicates.

### Chronic kinase perturbation

For chronic kinase perturbation, small molecule inhibitors (Supplementary Table 4) were applied to the cells at a concentration of 1 μM in parallel with TGFβ or mock treatment. The small molecule inhibitor was applied once per day for 5 days, after media change and 10 minutes before TGFβ stimulation, and collected in two replicates.

### Data preprocessing

All data were arcsinh transformed with a cofactor of 5 [2]. Any remaining debris or doublets were removed by gating on the DNA channels. For the time course and acute inhibition validation, the raw data was cleaned to remove cells that had spuriously high levels of certain signaling markers and transcription factors (pCREB, pSTAT5, pMEK1/2, pNFKB, Twist, Snail1 and Slug). An example between pCREB and pMEK1/2 is shown in Supplementary Figure 8A. The effect is seen only in the markers whose metal antibodies have similar masses (Supplementary Table 1), hence indicating that the high correlation could be experimental noise. Further uninformative cells that had low levels of all markers were removed. For this, cells were clustered using Phenograph [5] on a set of phenotypic markers and transcription factors (E-cadherin, Vimentin, CD24, CD44, βcatenin, Snail1, Slug and Twist). The clusters of cells with low levels of markers were discarded thereafter (Supplementary Figure 8B). The *junk* cells present in the data used for validation via acute inhibition (Figure 5 and Supplementary Figure 6) were also removed using Phenograph on the set of available phenotypic markers and transcription factors (E-cadherin, Vimentin, CD24, CD44, β-catenin, Snail1 and Slug).

### Assessing cellular heterogeneity

We quantified the proportion of cells that complete the transition (Figure 1C and Supplementary Figure 1, Supplementary Figure 2) by manually gating cells into various stages based on the expression levels of the canonical markers, E-cadherin and Vimentin. Cells with expression level of Vimentin < 2 were defined as epithelial cells, those with E-cadherin < 2.5 and Vimentin > 4 were defined as mesenchymal and rest of the cells as transitional. The same gates were used for all time-course data.

### Overview of computational methods to quantify edge dynamics

The computational methods developed in this paper are geared towards learning time-varying edge dynamics from static snapshot data. We study pairwise relationships as a function of time in cells undergoing the epithelial-to-mesenchymal transition. Studying such cell state dynamics from a single time point require computational techniques that can efficiently harness the rate variability within large samples of cells to capture the transient dynamics.

We develop information theoretic techniques to study edge relationships as a function of pseudo-time. These methods quantify the edge strength and describe time-varying edge shape. In particular, we develop:

1. *3D-DREVI (3D conditional Density Rescaled Visualization)* to visualize and characterize the relationship between a pair of molecules, *Y* and *Z*, along time *T*. For this, we compute the conditional density estimate 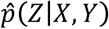 to capture the dependency of *Z* on *T* and *Y* and use it visualize the average expression of *Z* given *Y* and *T*.
2. *3D-DREMI* (3D conditional Density Resampled Estimate of Mutual Information) to quantify the strength of relationship of *Z* on both *T* and *Y* by computing the differential entropy of *Z* when conditioned on *T* and *Y*.
3. *TIDES (Trajectory Interpolated DREMI Scores)* to quantify the relationship between two molecules continuously along time. This involves computing 2D-DREMI on fixed-time slices in the 3D space to derive the time-varying strength of the relationship.

First, we use Wanderlust [6] to align cells along a one-dimensional EMT-trajectory, which we call EMT-time. We treat EMT-time (*T*) as the *X* variable and a pair of molecules as *Y* and *Z* variables in order to compute 3D-DREVI, 3D-DREMI and TIDES. Underlying all our methods is the estimation of the joint density 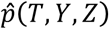, obtained using a fast heat-diffusion based kernel density estimate [7], which we extended to 3 dimensions. The methods are detailed as follows.

### Kernel Density Estimation

Kernel Density Estimation (KDE) is a data-driven approach for learning the underlying probability density function [8]. Given a set of points in 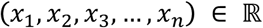, a kernel density estimate for the distribution of the points is given by,

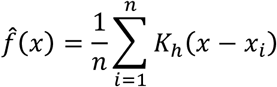

where, *K_h_* is the kernel function. A popular choice of kernel is the Gaussian kernel, given by,

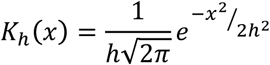

where, *h* is the bandwidth of the kernel. In higher dimensions, the kernel density estimate has the same form with points replaced by vectors.

#### Heat-Equation based KDE

A standard method for computing a kernel density estimate amounts to evaluating a kernel function, *K_h_*, at every data point and summing the result. However, this method can be computationally challenging for large data sets. Instead, we use a method based on heat diffusion [7], which has previously been used successfully in single cell data sets [9] for 2D-KDE. The method estimates the underlying distribution by modeling it as the spreading of heat governed by the heat equation (with delta functions at the data points as the initial condition). The intuition is that the fundamental solution to the heat equation, in an infinite domain with Dirac delta function as the initial condition, is a Gaussian function. Mathematically, the solution to

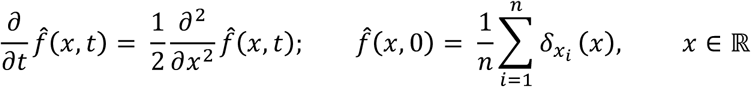

is given by,

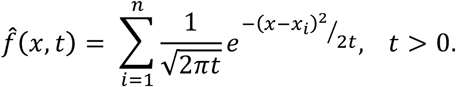

For practical purposes, we have finitely many data points, so we rely on the finite domain solution to heat equation approximation of the kernel density estimate. We enforce Neumann boundary conditions (derivative of the probability density function is 0 at the boundaries), which preserves the total probability mass (initial amount of heat) inside the boundary. Given the initial condition and the boundary conditions, the solution to the heat equation can be written as [10],

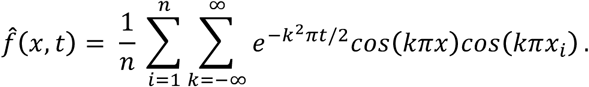

The solution can be efficiently computed using a fast Fourier transform (FFT) [11]. This results in an estimate of the underlying probability density function. For 1- and 2- dimensions, the bandwidth (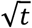) is obtained as a non-parametric solution to a fixed-point iteration [7, 11]. However, this method of obtaining bandwidth does not generalize beyond 2-dimensions [12] and becomes expensive to compute numerically. To generalize these ideas to higher dimensions, in this case 3-dimensions, we choose the bandwidth using Silverman’s rule of thumb [13],

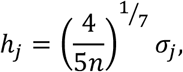

where *n* is the number of points, *σ_j_* is the standard deviation in *j^th^* direction.

#### Algorithm

The algorithm starts by binning the data into a histogram (with number of bins set by the user). This is already a rough estimate of the underlying probability density. Although fast to construct, a histogram is not smooth, over-fits the data and depends heavily on the size of the bin. However, the strength of the presented algorithm lies in the fact that the resulting histogram is treated as delta functions on equally spaced points and this is used as the initial condition for solving the heat equation. This reduces the sample space from the original data size to the number of bins, hence achieving a considerable gain in speed. Then we transform the data into the frequency domain using the discrete cosine transform (DCT), which can be implemented using FFT, applied onto this initial condition. This separates the signal present in the histogram into high frequency (noise) and low frequency (informative), thus allowing us to remove the noise and preserve meaningful information. The transform is then allowed to evolve for a time *t* (square of the bandwidth obtained using the rule of thumb), which is equivalent to multiplying by the exponential term (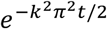) in the equation above, which is equivalent to low-pass filtering of the DCT. The resultant is then inverted (inverse-DCT) to obtain a smooth kernel density estimate, see Supplementary Figure 9. The method extends naturally to higher dimensions. Computing kernel density estimates using heat diffusion can be performed in *O*(*n* + *m* log*m*) ~ *O*(*n*) for *n* ≫ *m*, where *n* is the number of data points, *m* is the number of bins. A sketch of the algorithm is as follows [11],

1. Construct an equi-binned histogram
2. Transfer histogram into frequency domain via a discrete cosine transform
3. Evolve DCT (multiply DCT by 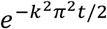, where *t* is the square of the bandwidth, *k* = 1,…, *m*, and *m* is the number of bins)
4. Inverse DCT for solution.

### 3D-DREVI

As with 2D-DREVI [9], the joint high-dimensional density estimate can reveal areas of the state space that are densely and sparsely occupied by cells. However, as dimensionality increases, the sparsity of the data, relative to the state space, has increasingly larger impact. Already in 3 dimensions, the joint density of variables *p*(*X*, *Y*, *Z*) is often not good at revealing the underlying relationship between *X*, *Y* and *Z* because the majority of cells may be within a restricted portion of the dynamic range. Therefore, to accentuate the dependencies between molecules, we consider the conditional relationship of *Z* given *X* and *Y*, thus capturing the dependencies across the full dynamic range [9]. To compute conditional density *p*(*Z|X*,*Y*), we normalize the joint density by the conditioning variables *X* and *Y*. Since it is difficult to visualize a 3-dimensional conditional density, we instead visualize the conditional mean of *Z* given *X* and *Y*, resulting in a 2D surface (Figure 3C).

#### Computing 3D-DREVI

We begin by computing a 3D kernel density estimate 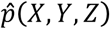 on a cubic mesh grid {*x_i_*, *y_j_*, *z_k_*, 0 < *i*, *j*, *k* < *m*, *m* is the number of bins using our heat equation based approach described above. Then each vector in the *z*-axis (corresponding to a fixed *X*- and *Y*-value, *X* = *x_i_*, *Y* = *y_j_*) is renormalized by the marginal density estimate of -*X* = *x_i_*, *Y* = *y_j_*,

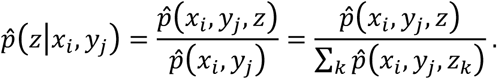

We thus obtain an estimate for the underlying conditional density *p*(*Z|X*,*Y*) on the cubic mesh grid.

#### Visualizing 3D-DREVI

Computing 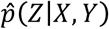 results in a 3-dimensional array, where each entry represents the value of the density estimate at a particular vertex on the 3D-mesh grid, making it difficult to visualize what is essentially a solid cube. Instead, we visualize a surface through the conditional mean of *Z* given *X* and *Y*. This incidentally is often the area of the highest conditional density. The conditional mean can be computed as follows,

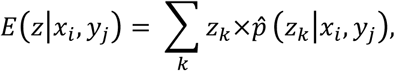

which results in a matrix where each entry corresponds to the average value of *Z* conditioned on the values of *X* and *Y*. This can be depicted as a surface plot on the *X* and *Y* mesh plane.

In this manuscript, we use 3D-DREVI to illustrate the relationship between two molecules along EMT-time (e.g. Figure 3D-E). We treat the Wanderlust derived EMT-time (*T*) as the *X* variable and the two molecules as *Y* and *Z* variables. We estimated the joint density at 2^21^ points (128 bins in each axis) and obtain the conditional mean of *Z* given *T* and *Y* as described above and render it as a red surface plot. Given a pair of molecules (*Y* and *Z*) and EMT-time, 50 cells from the right tail of the distribution of *Y* were discarded to obtain a well-populated dynamic range of Y, analogous to [9]. Finally, we remove wrinkles from the surface by smoothing the conditional mean using a linear sliding filter (of span 20 along both *T* and *Y* axes), using the *smooth* function in MATLAB.

### 3D-DREMI

Once 3D-DREVI is computed, we compute 3D-DREMI, an extension of DREMI [9], to quantify the strength of Z’s dependency on *X* and *Y*. Similarly to DREVI, we evaluate the strength of this dependency by re-weighing the contribution of each grid point uniformly thus taking the full dynamic range of the function into account [9].

#### Computing 3D-DREMI

Given three variables *X, Y* and *Z* (we typically assume that *X* and *Y* both influence *Z*), we quantify the dependence of *Z* on both *X* and *Y*. 3D-DREMI is defined as the mutual information on data that is sampled from the rescaled denoised-conditional density,

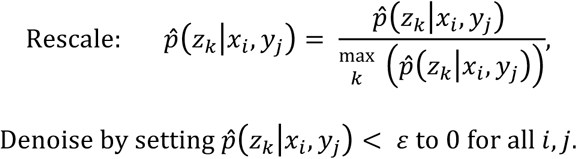

We measure the change in entropy of *Z* when conditioned on both *X* and *Y*, by computing the differential entropy between *Z* and *Z|X*,*Y*. That is, compute

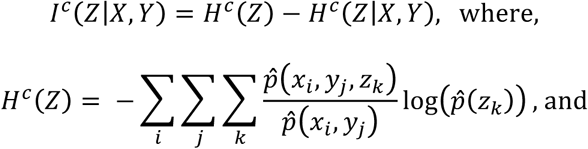

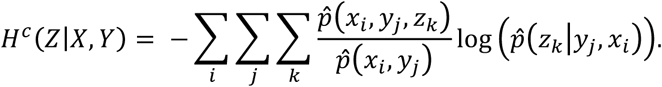

This is a natural extension of 2D-DREMI as detailed in [9]. By treating EMT-time (*T*) as the *X-*variable, we can assess the strength of relationship between *Y* and *Z* throughout EMT-time. Pairs with high 3D-DREMI scores have a strong relationship with each other and this relationship changes during the course of EMT-time. By ordering edges based on their 3D-DREMI score, we find candidate proteins that might be critical during EMT (Supplementary Table 3).

### TIDES

3D-DREMI quantifies the relationship between two molecules throughout EMT-time. However, it does not provide information about the strength of the relationship at a given EMT-time. We developed a method based on 2D-DREMI to evaluate how a relationship changes continuously with EMT-time. We call this method TIDES for Trajectory Interpolated DREMI Scores.

#### Computing TIDES

We start with the rescaled conditional density estimate of *Z* given *T* and *Y*, where we consider EMT-time (*T*), as the *X -*variable. This 3-dimensional density estimate is projected onto a slice along the *Y*-*Z* plane, resulting in the conditional dependence of *Z* on *Y* for various fixed values of *T*. The projections are obtained by linearly interpolating the 3D density estimate onto a 2-dimensional slice, {(*t_i_*, *y_j_*, *z_k_* : 0 < *j*,, *k* < *m*, and *i* is a fixed value, *m* is the number of bins}, along *Y* and *Z* direction, for which we use the “interp3” function in MATLAB. The resulting conditional density estimate is denoised at *ε* = 0.9 to eliminate the technical noise from measurement [9],

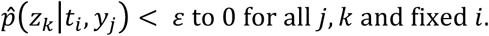

2D-DREMI computed on the slice quantifies the relationship at the fixed EMT-time *T*,

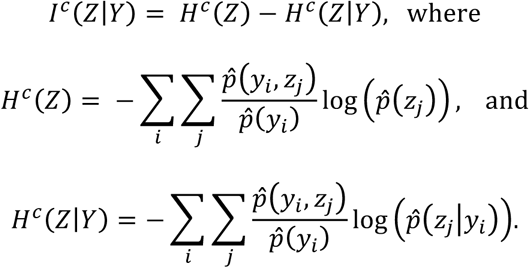

#### Visualizing TIDES

Repeatedly computing TIDES for several values along EMT-time allows continuous tracking of edge strength during the EMT transition, resulting in a TIDES curve. We compute TIDES values at 256 locations along EMT-time, which is twice the number of bins used to estimate the density. Once computed, the TIDES curves were smoothed using a Gaussian filter. For this, a Gaussian centered at each value of EMT-time (on which TIDES is computed) is used to estimate the weighted average at each location. Averaging the values results in a smooth TIDES curve. The weights are determined as follows,

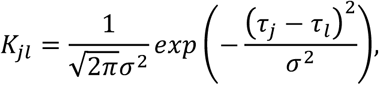

where, *τ_j_* is the TIDES value at EMT-time *j*, *τ_t_* is the mean TIDES value in the bin *l* and u is the bandwidth of the Gaussian chosen using Silverman’s rule of thumb [13]. The weighted average is then calculated as,

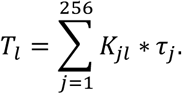

### Deriving Wanderlust pseudo-time

We used Wanderlust [6], a graph-based trajectory detection algorithm, to align the cells onto a one-dimensional axis representing the transition of cells from epithelial to mesenchymal phenotype. We call the resulting pseudo-time axis as EMT-time. EMT-time is normalized to 0-1, where epithelial cells are near 0 and mesenchymal cells near 1. We compute EMT-time by running Wanderlust on a set of phenotypic markers and transcription factors: E-cadherin, Vimentin, CD44, βcatenin, Snail1, and Slug. A shared nearest neighbor graph was constructed using *K* = 60 nearest neighbors and shared nearest neighbor (snn) = 20. The parameter *l* which is used to choose *l out of K* neighbors (to avoid short circuits) was set to K/5 = 12. The constructed trajectory is robust to these parameters (Supplementary Figure 10). The *start* point was set to the set of the cell with low E-cadherin and high Vimentin. In particular, the cell with maximum expression of Vimentin from the set of cells whose expression of E-cadherin < 1.5 and Vimentin > 4.5 was chosen as the *start* point. The resulting trajectory was then inverted. The number of graphs (over which the result of the algorithm is averaged) was set to 5.

Once generated, we study the expression of various markers along EMT-time (e.g. Figure 2B-C, Supplementary Figure 3A-C). The marker trends were generated by first partitioning EMT-time into 256 equally spaced bins, by dividing the range of the Wanderlust score by 256. Then the weighted average of the marker using a Gaussian filter centered at the bin is computed, as detailed in [14]. The weights are calculated as follows,

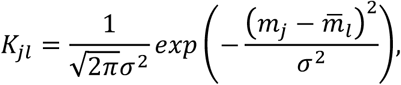

where, *m_j_* is the marker expression of cell *j*, 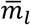 is the mean marker expression value in the bin *l* and *σ* is the bandwidth of the Gaussian chosen using Silverman’s rule of thumb [13]. The weighted average is then calculated as,

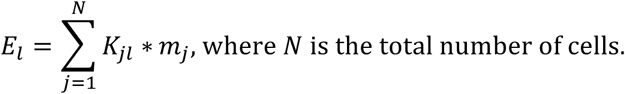

#### Consistency of marker trends along EMT-time across replicates

We demonstrate that the marker trends are consistent across replicates (Supplementary Figure 3D). For a given day, the expression of a marker along EMT-time from one replicate was cross-correlated with the expression of the same marker along EMT-time from another replicate. This was repeated for all the markers and average correlation was computed. The computation was done for all 3 replicates from Day 2, 3 and 4. The markers used were: pCREB, pSTAT5, pFAK, pMEK1/2, Twist, cmyc, Snail1, pNFKB, pP38, pAMPK, pAKT, pERK1/2, Slug, Cyclinb1, cah, pGSK3β, pSMAD1/5, CD44, Vimentin, pSMAD2/3, β-catenin, pMARCK, CD24, pPLCγ2, pPH3, pS6, E-cadherin, ccasp, pSTAT3, pRb, Survivin.

#### Consistency of marker trends along EMT-time across days

We also find that the marker trends are consistent across days (Supplementary Figure 3E). The expression of a marker along EMT-time on replicate 1 of Day 2 was correlated with the expression of the same marker along EMT-time on replicate 1 of Day 3. This was repeated for all the markers and average correlation was computed. The same is done to compare replicate 1 of Day 2 against Day 4 and replicate 1 of Day 3 against Day 4. The final result is rendered as a heat-map. Similar heat-maps were generated for replicates 2 and 3. The markers used were: pCREB, pSTAT5, pFAK, pMEK1/2, Twist, cmyc, Snail1, pNFκB, pP38, pAMPK, pAKT, pERK1/2, Slug, Cyclinb1, cah, pGSK3β, pSMAD1/5, CD44, Vimentin, pSMAD2/3, β-catenin, pMARCK, CD24, pPLCγ2, pPH3, pS6, E-cadherin, ccasp, pSTAT3, pRb, Survivin.

#### Consistency of signaling controlled for EMT-time

We demonstrat that signaling is similar across days when controlled for EMT-time (Figure 3A-B, Supplementary Figure 4A). Cells from days 2, 3 and 4 were divided into four groups based on EMT-time: cells with EMT-time < 0.25 (Group-1), between 0.25 and 0.5 (Group-2), between 0.5 and 0.75 (Group-3) and greater than 0.75 (Group-4). DREMI on all pairs of signaling molecules in each of the groups was computed. Then the DREMI scores from a group was correlated with the DREMI scores from the same group on a different day, and the final result is rendered as a heat map. The markers used are: pCREB, pSTAT5, pFAK, pMEK1/2, Twist, cmyc, Snail1, pNFκB, pP38, pAMPK, pAKT, pERK1/2, Slug, Cyclinb1, pGSK3β, pSMAD1/5, pSMAD2/3, β-catenin, cah, pMARCK, pPLCγ2, pPH3, pS6, pSTAT3 and pRb.

### Validating short-term drug inhibition

We used short-term drug inhibition to validate rewiring suggested by TIDES. For a given pair of molecules *X* and *Y*, TIDES (*X* → *Y*) quantifies the strength of the statistical relationship between *X* and *Y* continuously along EMT-time. We assume that inhibiting *X* or some molecule immediately upstream of *X* should have higher impact on *Y* in the region where the TIDES score is high, and analogously the impact should lower in regions of lower TIDES scores. To validate TIDES, we compute TIDES curve of *X* − *Y* and cross-correlate it with the impact curve of *Y*, defined as the expression of *Y* under control minus the expression of *Y* under treatment. A high correlation would indicate that TIDES correctly predicts the regions of strong/weak relationship. For analysis, both the curves were normalized to -1 to 1 and cross-correlated on the EMT-time axis. The shift which provided the maximum cross-correlation was chosen and the Pearson correlation value was reported.

### Validating long-term drug inhibition

We defined manual gates, based on the levels of E-cadherin and Vimentin, for computing the fraction of mesenchymal cells to validate the impact of long-term drug perturbation on EMT (Supplementary Figure 7). The gates used are: (1) TGFβ-receptor, MEK and AKT inhibition: E-cadherin < 3, vimentin > 4. (2) AMPK-perturbation: Ecadherin < 3 and Vimentin > 3.5. (3) WNT-inhibition: E-cadherin < 3 and Vimentin > 4 (Replicate 1), and E-cadherin < 3.5 and Vimentin > 4 (Replicate 2). Our predictions are validated across replicates.

### Runtime Analysis

We performed runtime analysis of our methods. We first assessed how our method scales with size of the data. Since heat-diffusion based kernel density estimation starts off by computing the histogram of the data (Supplementary Figure 9), we fixed the number of bins to 128. For randomly generated data sets from uniform distribution, with sample size ranging from 5000 to 50000 (3 features for each data-point), the heat-diffusion based method computes KDE within 1 second, Supplementary Figure 11A. The runtime is uniform across a range of data size because the algorithm is less dependent on the data size and more on bin size, which was kept constant here. Second, we studied the run time of our method against the number of bins in the initial histogram. We fixed the size of the data set to 5000 points (each with 3 features) and altered the number of bins in the initial construction of the histogram. For up to 256 bins in each direction (density estimated at 2^24^ points), the heat-diffusion based method computes KDE within 10 seconds (Supplementary Figure 11B). We compared the runtime of our method to an alternative approach [15]. We used the code available at http://www.ics.uci.edu/~ihler/code/kde.html. As shown in Supplementary Figure 11C, our method scales better than the alternative against the number of bins of histogram. Using the heat-diffusion based KDE, 3D-DREVI and 3D-DREMI can be computed within 25 seconds for 128 bins (Supplementary Figure 11 (D)-(E)). Similarly, TIDES can be computed in less than 5 minutes (Supplementary Figure 11F). For all of these experiments, since the method depends mostly on the number of bins for the histogram, only an example pair of edges (pS6 -> pGSK3β) along EMT-time was chosen for three replicates from Day 3 (unless stated otherwise) and the average runtime was computed.

### Software Availability

Our software and computational methods are available at https://github.com/roshan9128/tides.

---

**Supplementary Figure 1.**
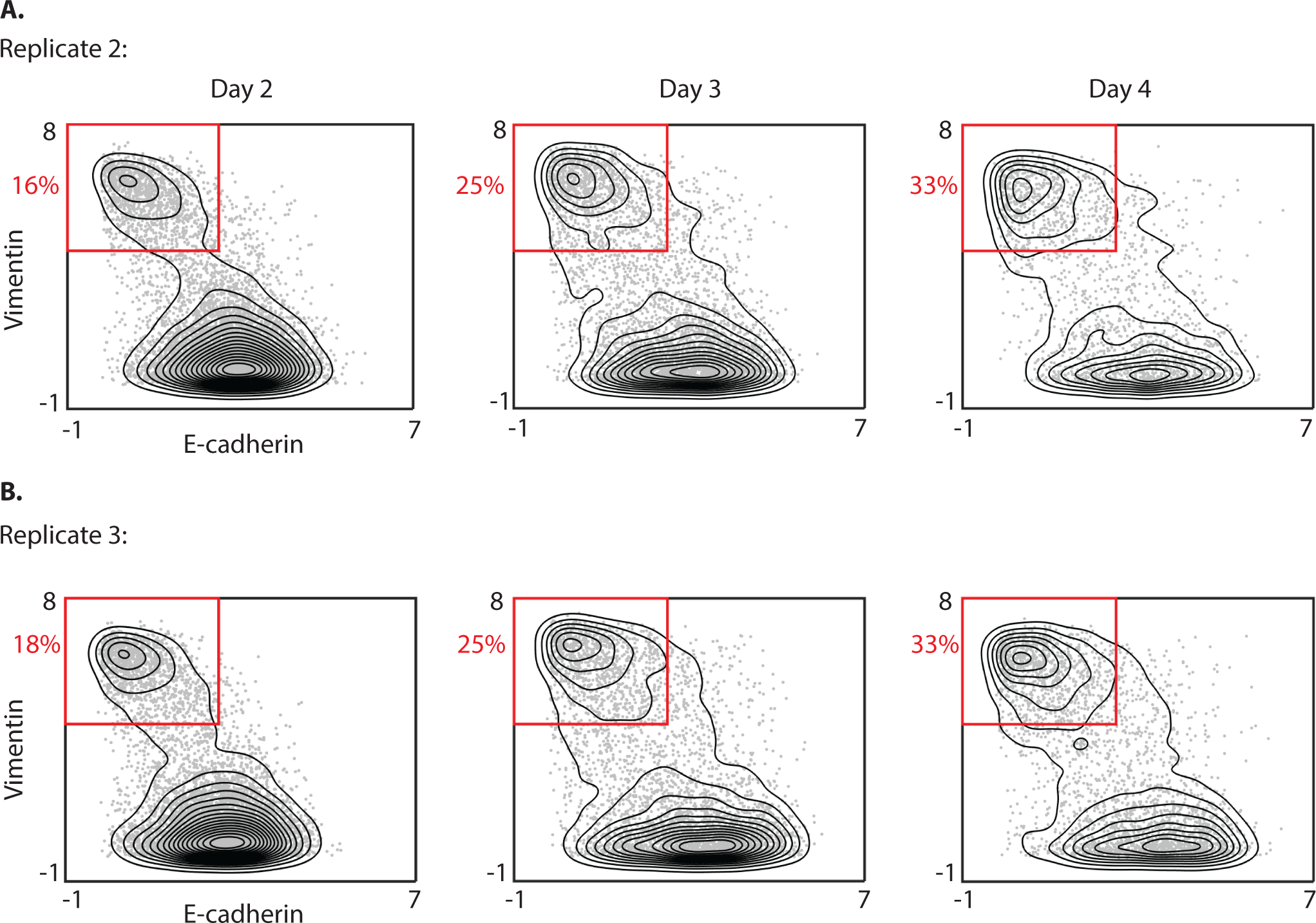
TGFβ treatment reproducibly induces EMT: (A-B) Contour plots of Vimentin and Ecadherin after 2-4 days of TGFβ stimulation; biological replicates for main Figure 1C. Replicates confirm a shift in density of cells from epithelial to mesenchymal phenotype with time and illustrate a continuum of cells in transition on days 2-4.

**Supplementary Figure 2.**
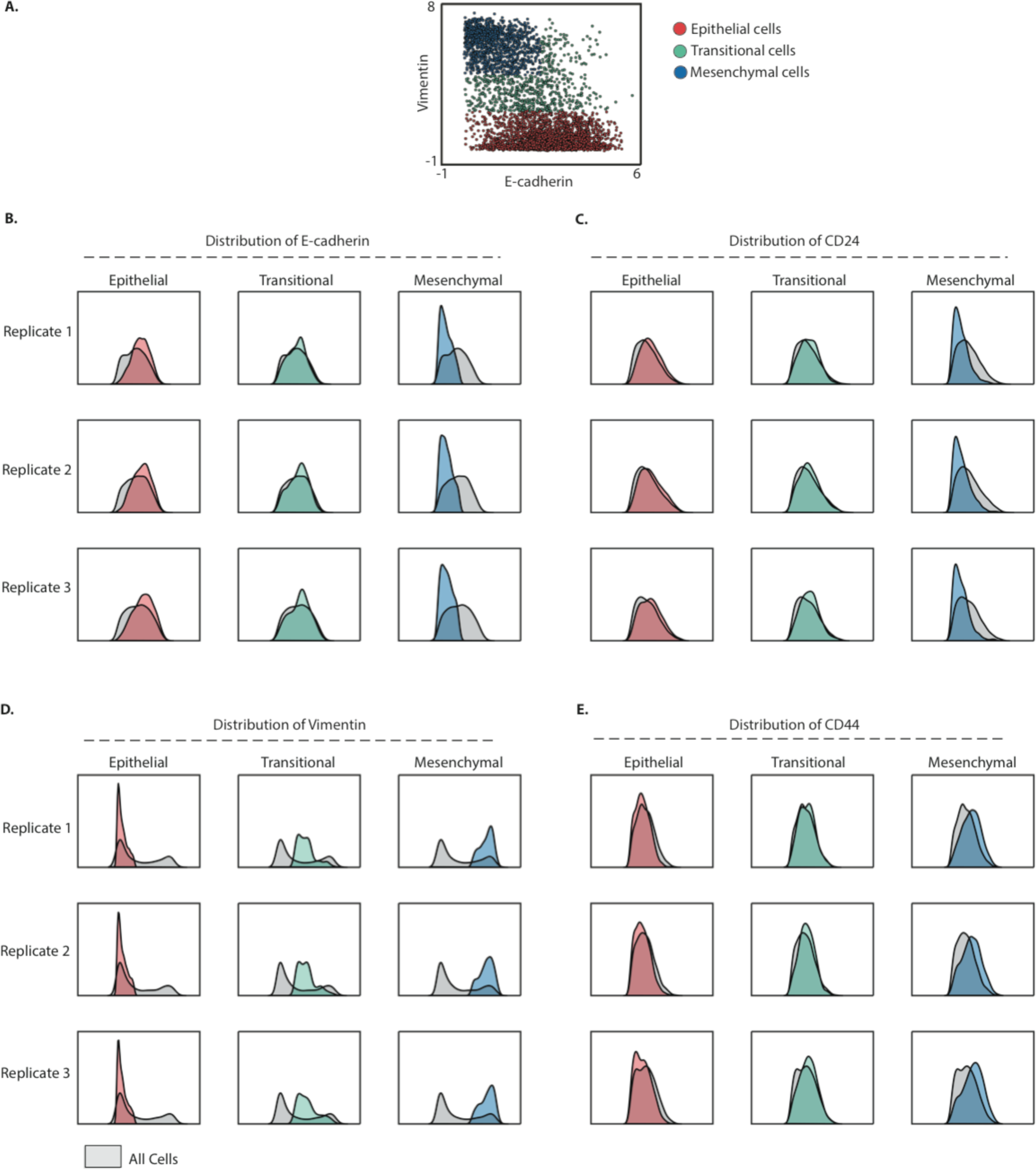
EMT characteristics are consistent across replicates: (A) Scatterplot where each point is a cell treated with TGFβ for 3 days. The cells are divided into three distinct categories: Epithelial, Transitional and Mesenchymal (see Methods). (B)-(E) A distribution of marker levels is shown for the three categories. Expression of E-cadherin (B) and CD24 (C) is high in epithelial cells, decreases in transitional cells, and is much lower in mesenchymal cells, consistently across replicates. Expression of Vimentin (D) and CD44 (E) is low in epithelial cells, increases in the transitional cells, and is higher in the mesenchymal cells, consistently across replicates.

**Supplementary Figure 3.**
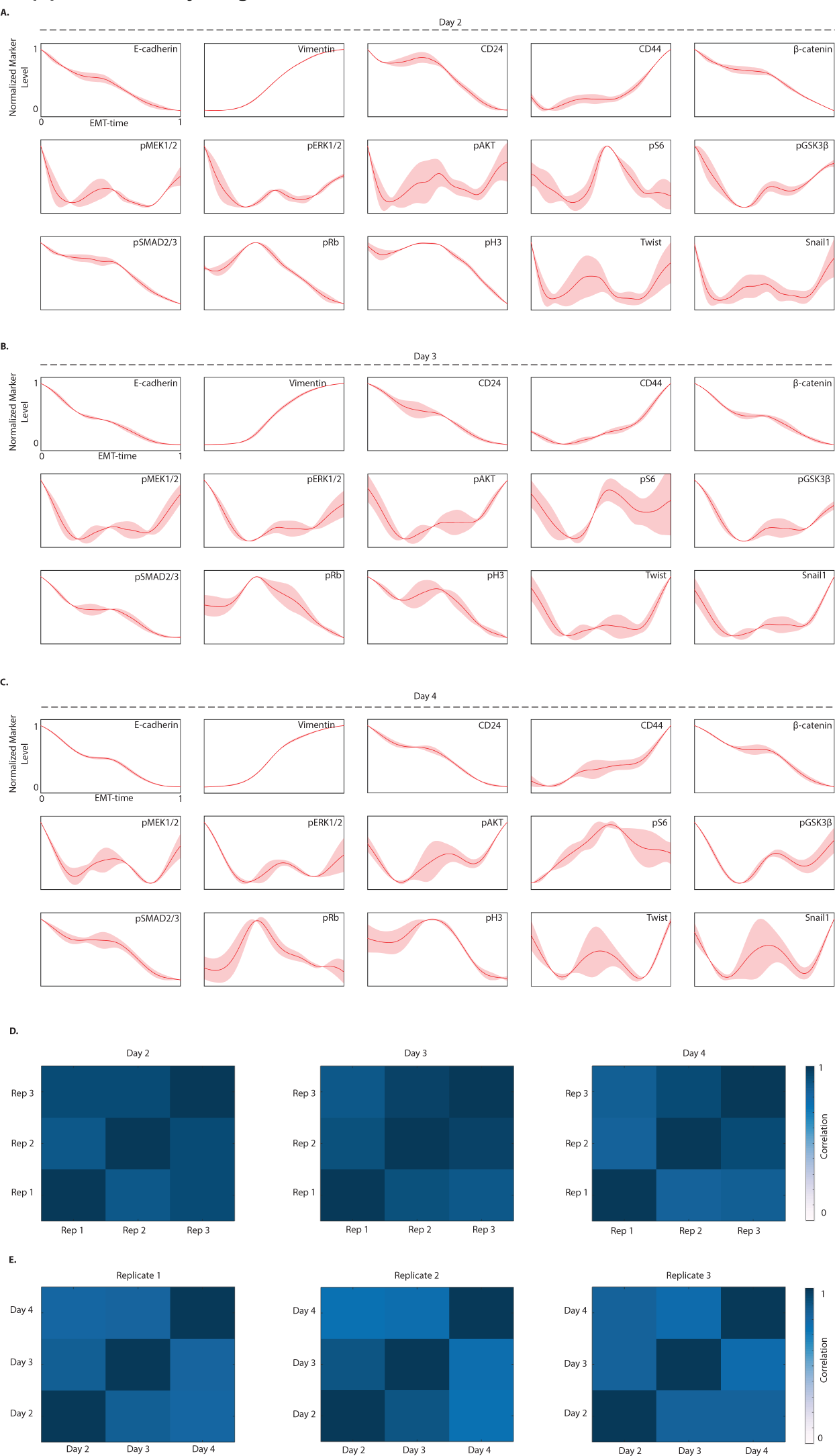
A spectrum of marker trends along EMT-time are seen consistently across replicates: (A)-(C) Plots show the expression of various markers along Wanderlust generated EMT-time in the cells treated with TGFβ on Day 2, 3 and 4 respectively. Smoothing was performed by a sliding-window Gaussian filter. The shaded region around each curve indicates one standard deviation across replicates indicating consistency. (D) Plot showing the average cross-correlation of marker expression along EMT-time across replicates. For a given marker, the expression along EMT-time is cross-correlated across replicates. The average correlation over the set of markers is rendered as a heat map. (E) Average cross-correlation of marker expression along EMT-time is similar across the different days within each replicate.

**Supplementary Figure 4.**
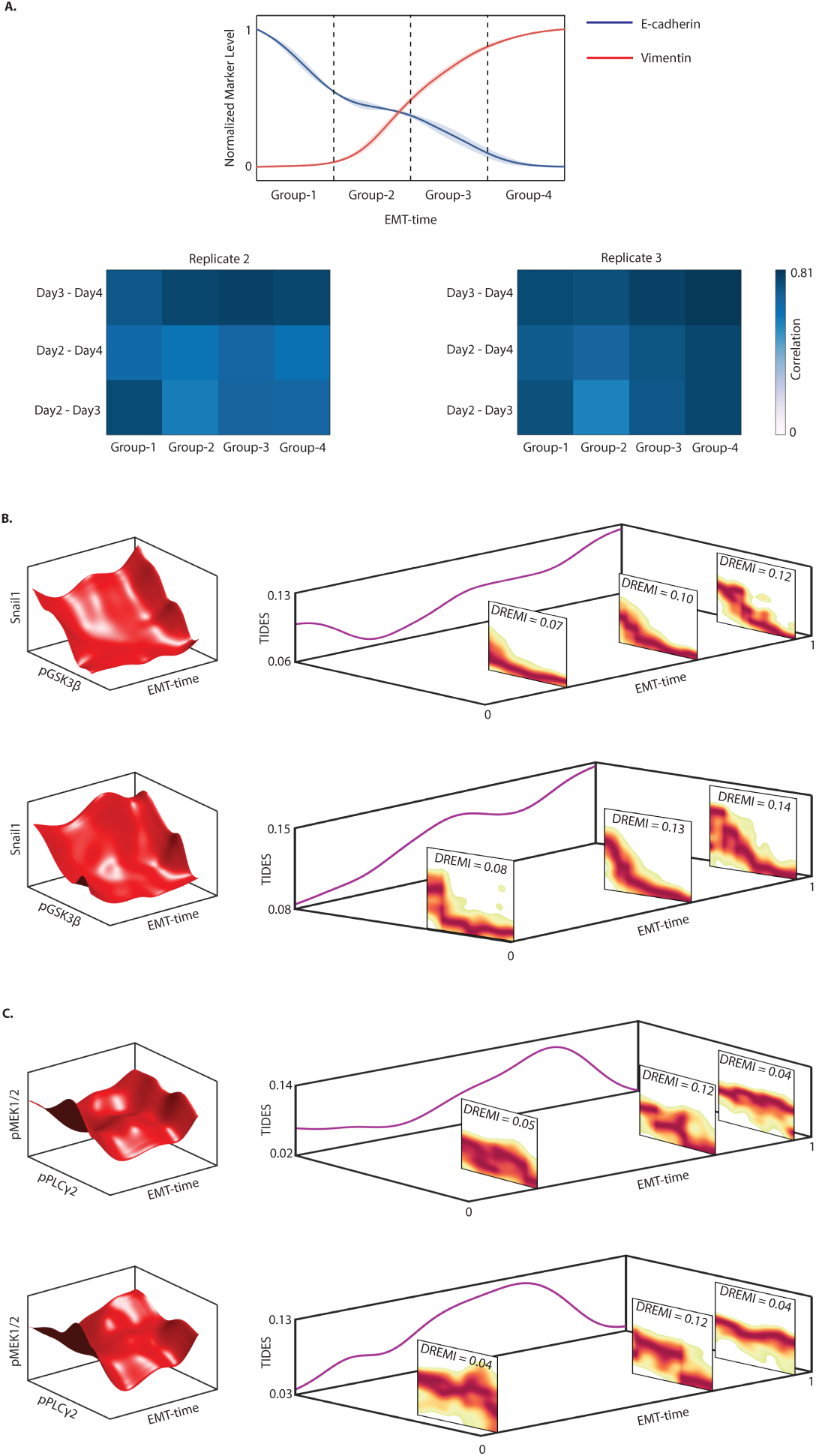
Signaling relationships along EMT-time in replicates: (A) TGFβ-treated cells from Days 2, 3 and 4 are binned into four groups along EMT-time. DREMI score between all pairs of signaling molecules is computed in each group. Heat map shows the correlation of the DREMI scores for each group across days. Average correlation is 0.68 (Replicate-2) and 0.73 (Replicate-3). (B) Dynamics of the relationship between pGSK3β and Snail1, similar to main Figure 3D across biological replicates. 3D-DREVI depicts the typical expression of Snail1 conditioned on pGSK3β and EMT-time. The modulation in the relationship is visualized by the 2D-DREVI slices along EMT-time and quantified the TIDES curve (purple curve) shown along the z-axis. (C) Dynamics of the relationship between pPLCγ2 and pMEK1/2 similar to Figure 3E across biological replicates.

**Supplementary Figure 5.**
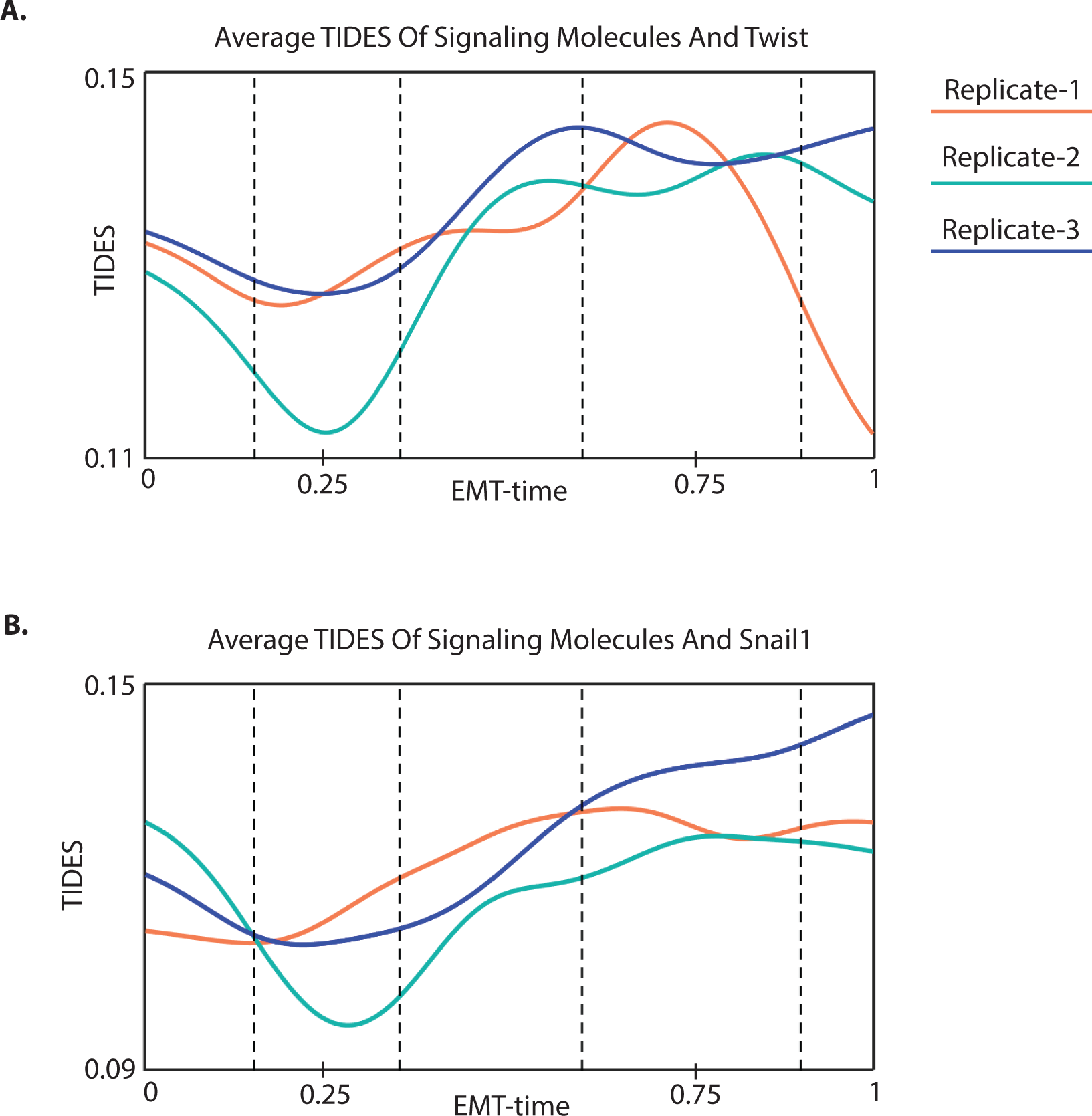
Information transfer during EMT across transcription factors: Average TIDES curve of the relationship between several molecules (pCREB, pSTAT5, pFAK, pMEK1/2, pNFκB, pP38, pAMPK, pAKT, pERK1/2, pGSK3β, pSMAD1/5, pSMAD2/3, β-catenin, CAH IV, pMARCK, pPLCγ2, pS6, pSTAT3) and Snail1 (B) and Twist (C), across three replicates for Day 3. Similar to Slug in main Figure 4, the curves start rising steadily at near EMT-time ~ 0.25, and peak near EMT-time ~ 0.75.

**Supplementary Figure 6.**
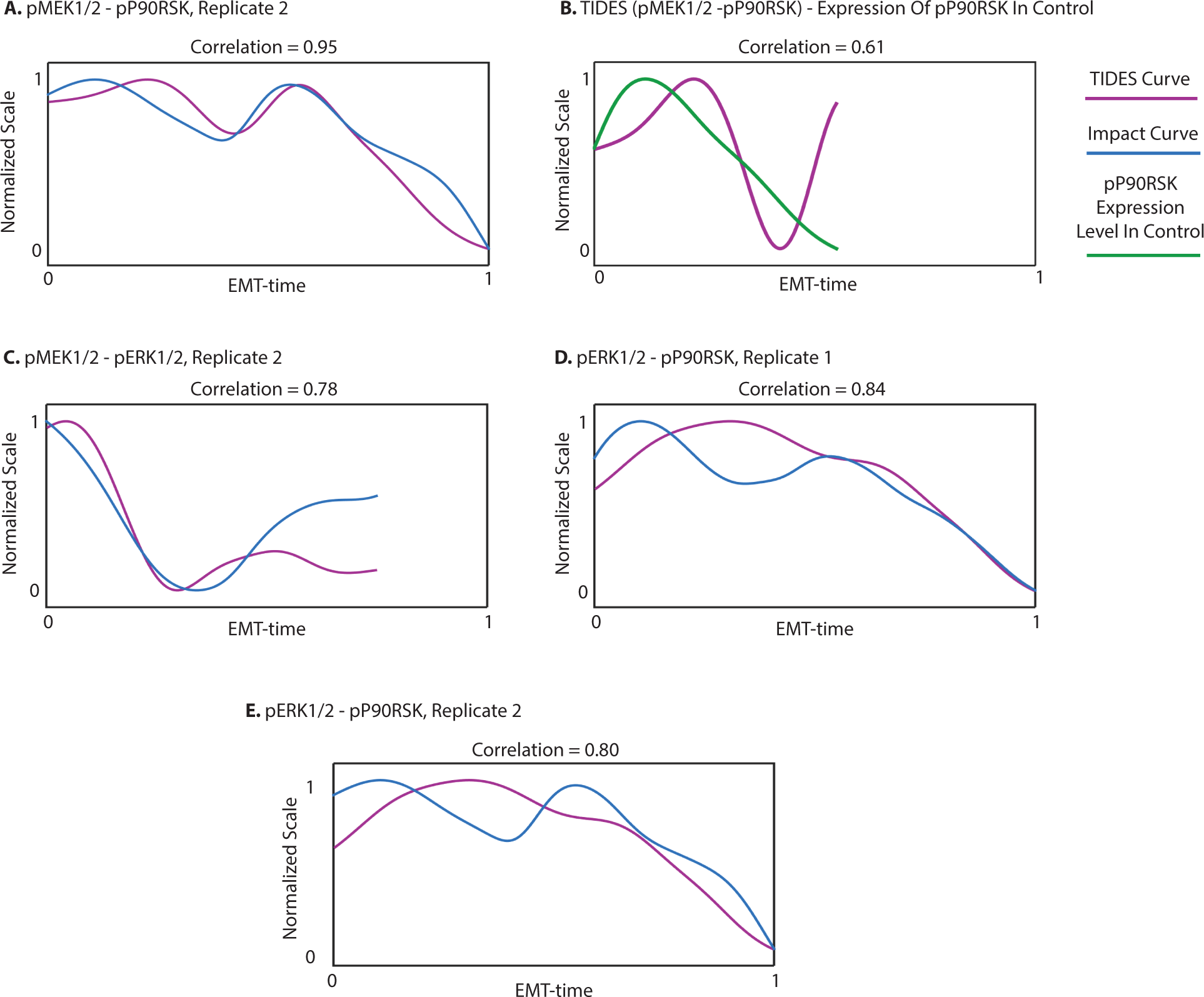
Validation of TIDES via short-term drug inhibition for direct and indirect edges in replicates: (A) Cross-correlation of TIDES curve between pMEK1/2-pP90RSK with the impact curve of pP90RSK results in a high correlation. This is a biological replicate of main Figure 5A. (B) Cross-correlation of TIDES curve between pMEK1/2-pP90RSK with the expression level of pP90RSK under control. Lower correlation than in (A) indicates that TIDES does not trivially follow the levels of pP90RSK. (C) Biological replicate of Figure 5B; cross-correlating TIDES curve between pMEK1/2-pERK1/2 with the impact curve of pERK1/2 results in a high correlation. (D)-(E) Cross-correlation of pERK1/2- pP90RSK TIDES curve and pP90RSK impact curve under MEK-inhibition is 0.84 and 0.80 across two replicates.

**Supplementary Figure 7.**
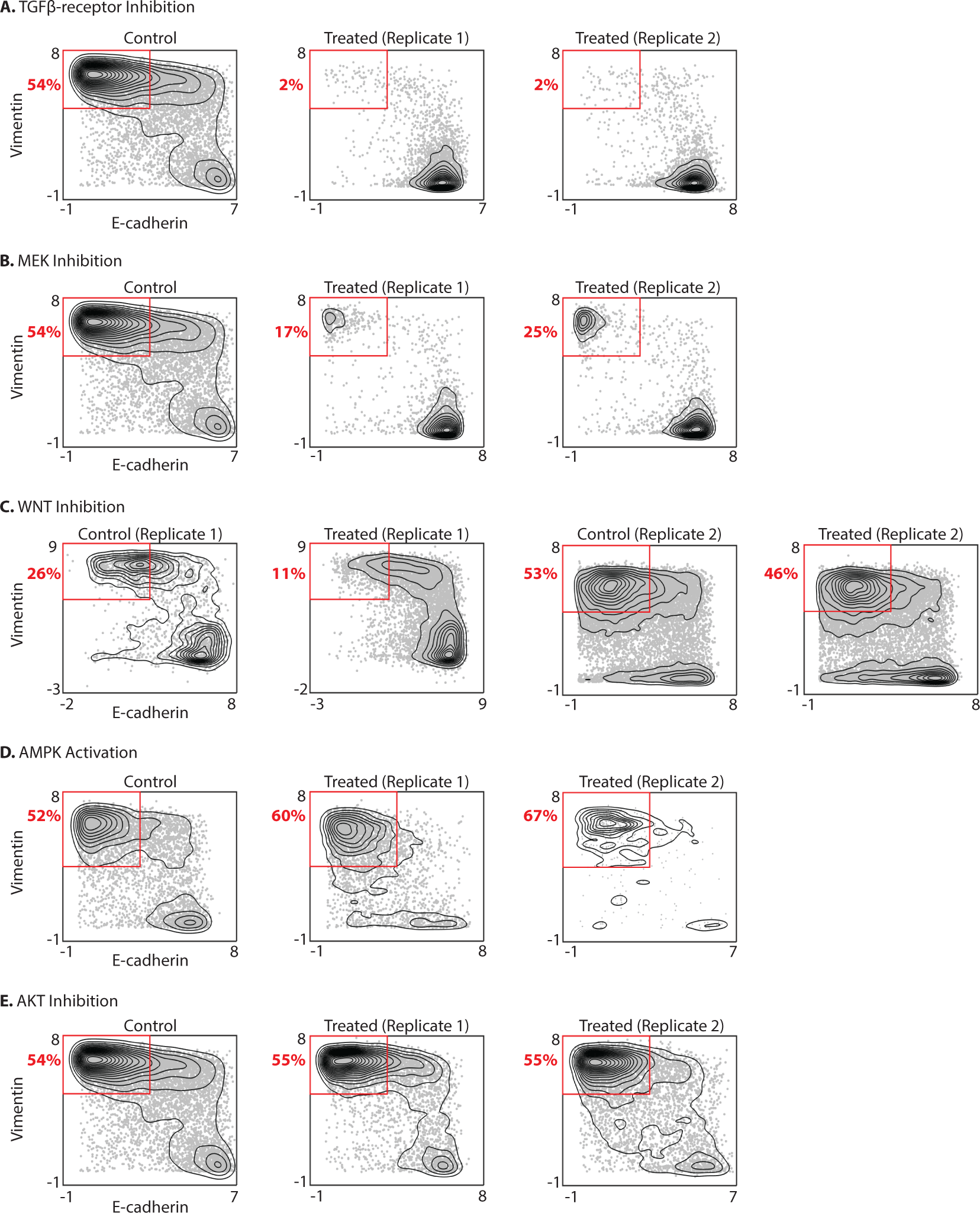
Validation of critical edges for EMT via long-term drug inhibition in replicates: (A)-(E) Shown are contour plots of cells treated with TGFβ (Control) and with TGFβ plus a chronic drug perturbation of the stated molecule for 5 Days, across biological replicates. Results of replicate 1 were shown as bar plots in Figure 6. Inhibition of TGFβ-receptor (A), MEK (B) and WNT (C) cause a substantial decrease in the fraction of cells that complete transition, while activation of AMPK (D) increases the proportion of cells that complete transition. AKT (E) on the other hand does not seem to impact the transition.

**Supplementary Figure 8.**
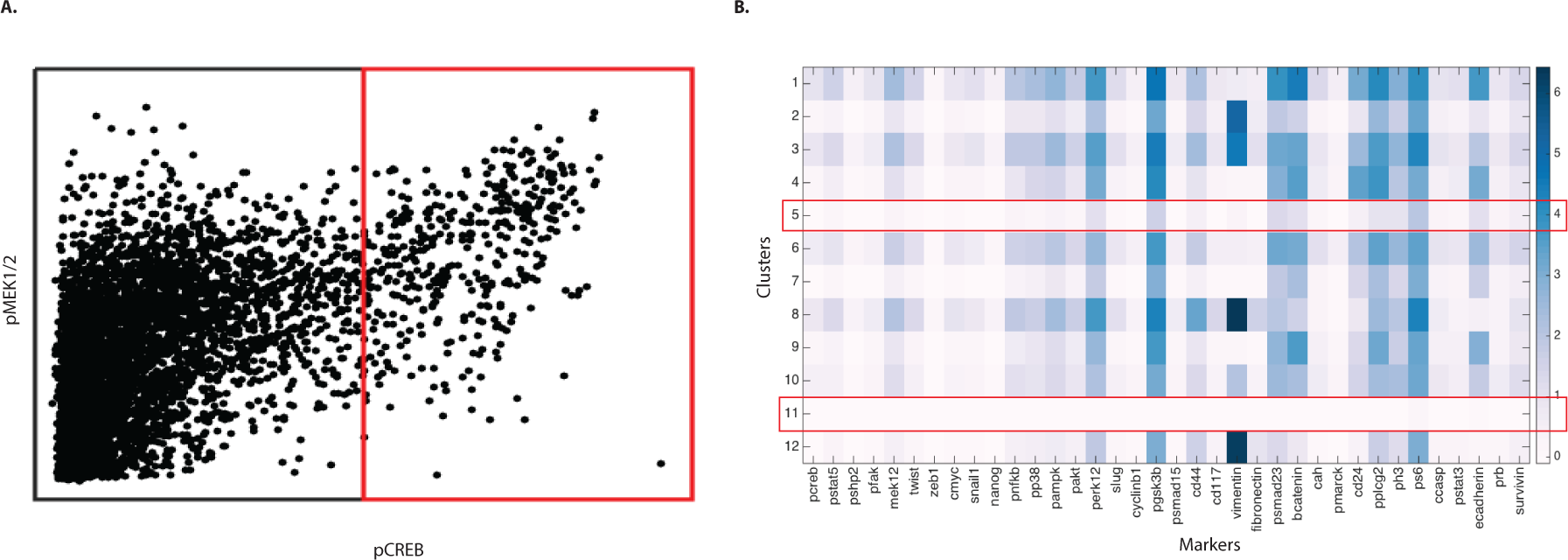
Data clean-up: (A). Scatterplot showing the relationship between pCREB and pMEK1/2 on Day 3 (shown is replicate 1). A spurious relationship between pCREB and pMEK1/2 at high pCREB values is seen. These events were manually gated out from time course and acute inhibition validation data sets. (B) Shown are heat maps of the expression of markers on various clusters obtained using Phenograph [1] on a set of phenotypic markers and transcription factors. The data shown is from Day 3 (replicate 1). The cells comprising the clusters with low expression of markers (such events are found in most mass cytometry experiments) were removed (indicated by red rectangles) from further analysis.

**Supplementary Figure 9.**
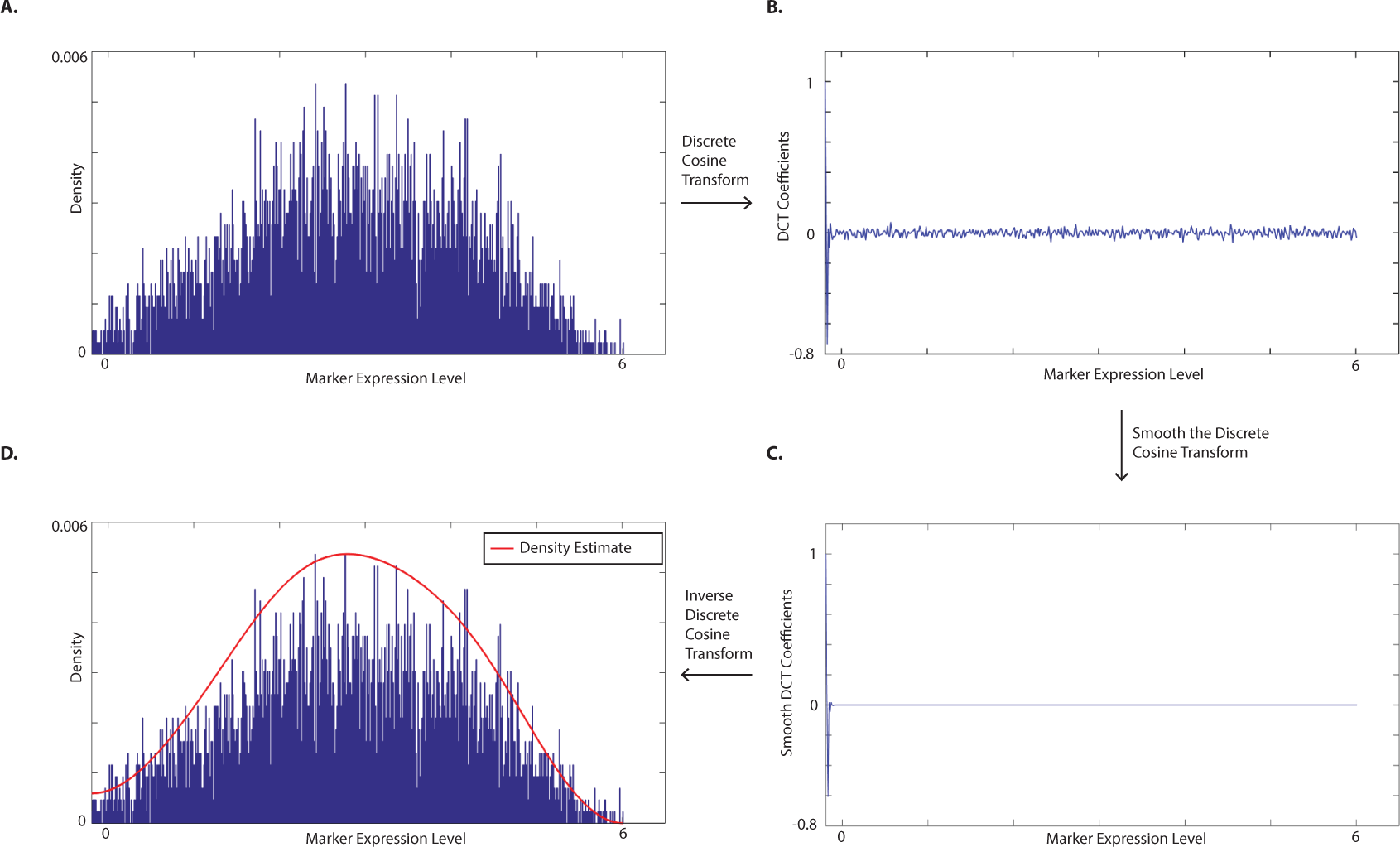
Computing Kernel Density Estimate: (Α) Plot shows histogram of a randomly chosen marker on Day 3. Constructing the histogram of the data is the first step in computing kernel density estimate. The histogram represents the initial condition for solving the heat equation. (Β) The histogram is then transformed into frequency domain by the Discrete Cosine Transform (DCT). (C) A low-pass filter smooths the DCT by removing the noisy parts. This is obtained by multiplying the DCT by an exponentially decaying term (exp(-k^2^π^2^t/2), see Methods) or in other words, evolving the initial condition in time. (D) The smooth density estimate is derived by inverting the smoothed-DCT.

**Supplementary Figure 10.**
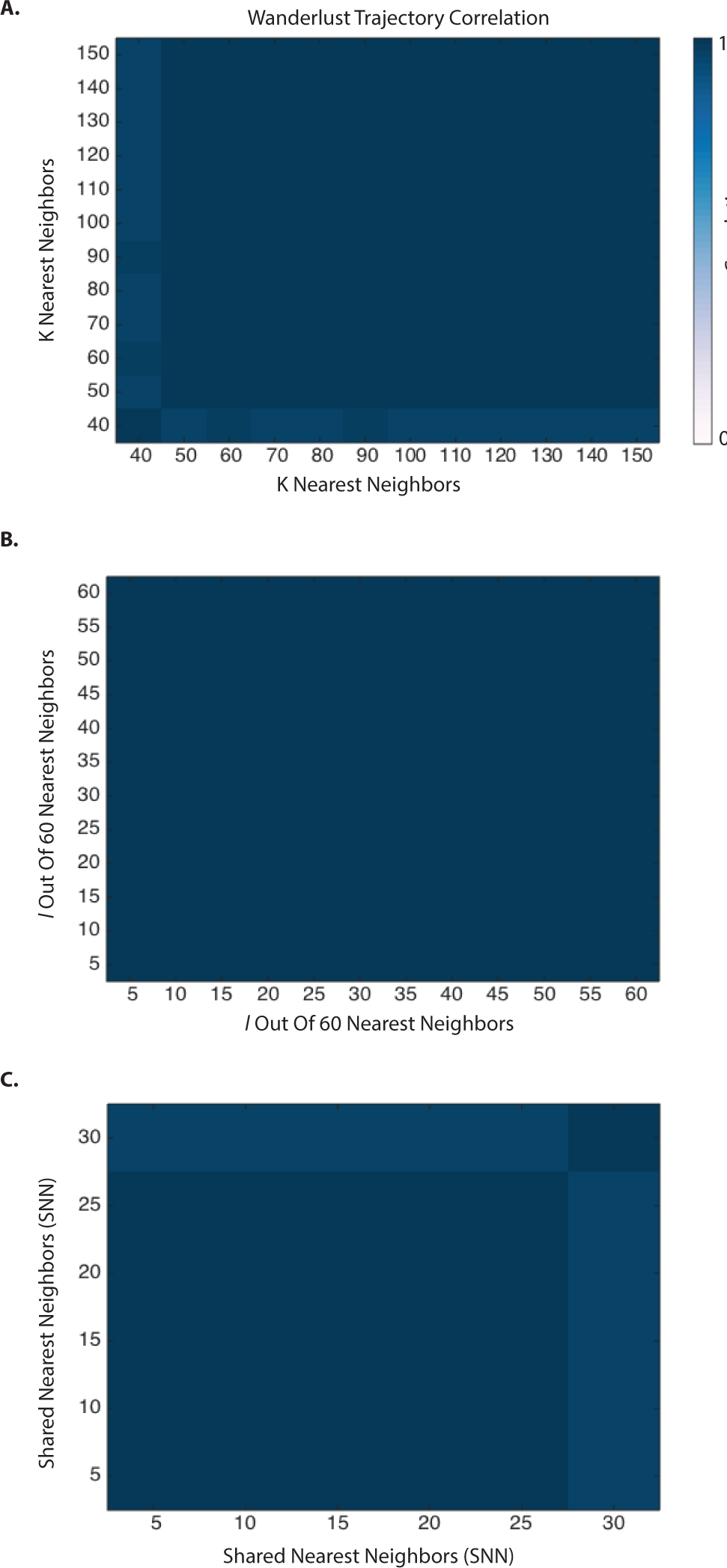
Robustness of Wanderlust generated EMT-time: (A) Heat-map shows the correlation between trajectories generated for various values of K-nearest neighbors. The shared nearest neighbor (snn) parameter was fixed at 20, while the *l* parameter (to avoid short circuits in the graph) was fixed at 12. (B) Heat-map shows the correlation between trajectories generated for various values of *l* parameter. K was fixed at 60 and snn was fixed at 20. (C) Heat-map shows the correlation between trajectories generated for various values of snn. K was fixed at 60 and *l* was fixed at 12. The results shown are for data on Day 3 (replicate 1), and holds true for all of our data.

**Supplementary Figure 11.**
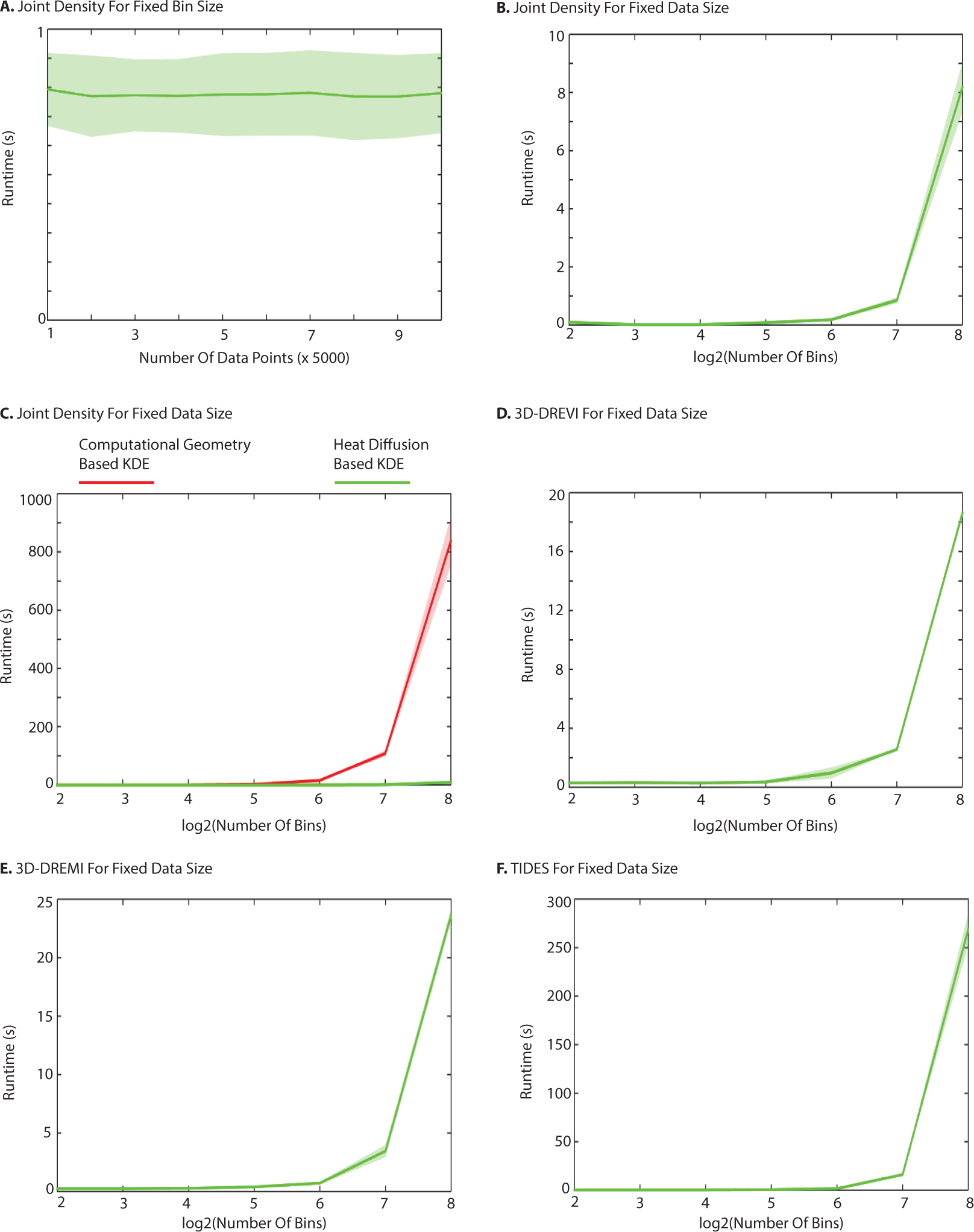
Runtime Analysis: (A) Runtime for computing heat-diffusion based KDE against data size. The data is uniformly generated with 3 features. The number of bins used to construct the initial histogram was fixed to 128 in each of the three directions. The green line shows the mean while shaded region shows one standard deviation for 100 iterations. (B) Runtime against the log of number of histogram bins (in each of the three directions) for 5000 uniformly generated points. Computing density estimates on 2^24^ points takes less than 10 seconds. (C) Runtime comparison of our method against an alternate (based on computational geometry) [2] for computing three-dimensional KDE. The shaded region shows the standard deviation across three replicates from the time-course data Day 3 for an edge (pS6-pGSK3β) and EMT-time. Our method can compute 3D density estimates at 2^24^ points in less than 10 seconds while the alternative takes more than 10 minutes. (D)-(F) Plots show runtime of heat-diffusion based method against log of number of bins in computing 3D-DREVI, 3D-DREMI and TIDES respectively. The shaded region shows the standard deviation in runtime across three replicates of data from Day 3 for the edge used in (C), and the middle line shows the mean runtime.

## Supplementary Tables

**Table 1:**
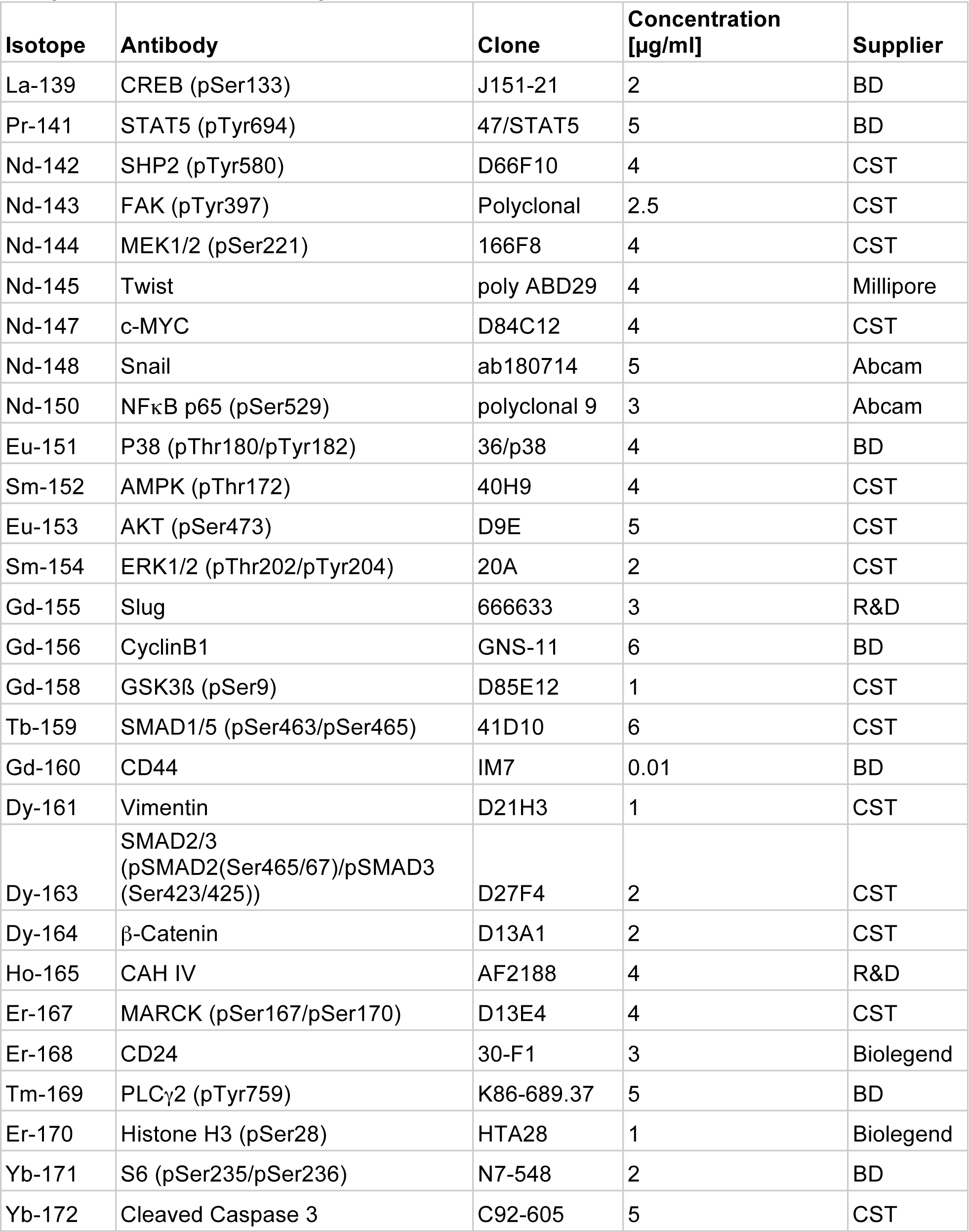
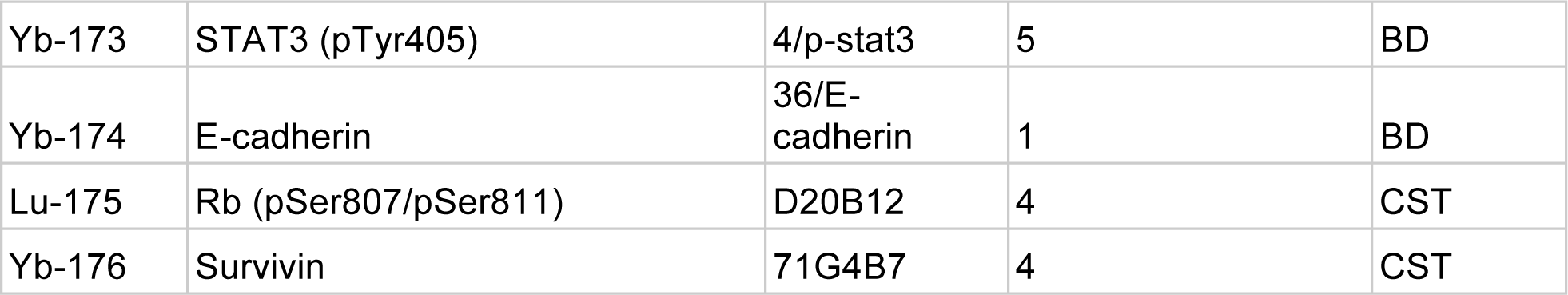
List of molecules and antibodies used for all mass cytometry analysis except the acute inhibition experiment.

**Table 2:**
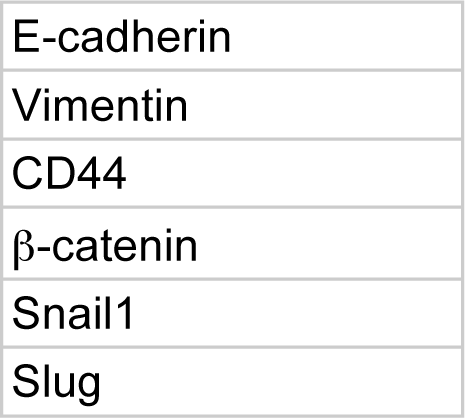
List of molecules used to construct wanderlust pseudo-time:

**Table 3:** List of small molecules used for chronic perturbation.

| X-molecule | Y-molecule | Average score |
| --- | --- | --- |
| pSMAD2/3 | β-catenin | 0.967 |
| pAMPK | β-catenin | 0.891 |
| pERK1/2 | β-catenin | 0.886 |
| pGSK3β | β-catenin | 0.885 |
| pGSK3β | pERK1/2 | 0.874 |
| pMEK1/2 | β-catenin | 0.866 |

**Table 4:** List of small molecules used for chronic perturbation.

| Target | Small molecule |
| --- | --- |
| TGFβ-R1 | SB431542 |
| MEK | PD318088 |
| WNT | XAV-939 |
| AMPK | Phenformin |
| AKT | PHT427 |

